# Sex-specific human electromechanical multiscale in-silico models for virtual therapy evaluation

**DOI:** 10.1101/2025.03.14.643310

**Authors:** Maxx Holmes, Zhinuo Jenny Wang, Ruben Doste, Julia Camps, Hector Martinez-Navarro, Hannah Smith, Jakub Tomek, Blanca Rodriguez

## Abstract

**Background and Aim:** Women are significantly under-represented in cardiovascular research and in the evaluation of treatment safety and efficacy, leading to poorer patient outcomes. Quantification and investigation of sex differences in human electromechanical function and underlying mechanisms is crucial. To address this, we present sex-specific human cellular and biventricular electromechanical models for mechanistic investigations into sex-differences in therapy evaluation through simulations.

**Methods:** Protein genomic expression data from healthy human myocytes were used to calibrate sex-specific models of human cellular electrophysiology, subsequently integrated in biventricular electromechanical models with male and female anatomies. A validation, verification and uncertainty evaluation were implemented at the cellular and biventricular level, including validation using sex-specific datasets from randomised controlled trials for Dofetilide and Verapamil, with known sex-differences. Ionic mechanisms underlying sex-differences in drug response were mechanistically investigated.

**Results:** Sex-specific electromechanical models recapitulate sex-differences from ionic currents to ECG biomarkers including QTc interval (Male: 312ms; Female: 339ms; 9% difference), T-wave amplitude (6-9% difference) and ST-steepness through electrophysiological changes alone. Sex-specific simulations demonstrate both ECG biomarkers and mechanical biomarkers (Female LVEF: 68%, Male LVEF: 50%) within healthy ranges in clinical data for male and female population in the UK Biobank (n= 806, 46% Male). ECGs sex-differences are primarily explained by ionic currents, whereas mechanical sex-differences are driven by anatomical differences, and secondarily through more robust calcium function in females. Under Dofetilide, simulations show exacerbated QT prolongation in women compared to men (54-78% increase in effect), and a T-wave amplitude decrease in males (up to 0.25 mV), consistent with clinical data. This is explained in simulations by lower repolarisation reserve in women (due to low potassium and high calcium currents) than men. Verapamil shows no effect on simulated QT in either sex, and divergent T-wave modulation (increased amplitude in females, decreased in males) consistent with clinical trial data. Simulations identify enhanced contractile reservoir in female compared to male, with lesser decreases to ejection fraction with calcium current block.

**Conclusion:** Simulations using novel, sex-specific cellular and biventricular electromechanical models reveal the primary role of ionic currents sex-differences in ECG and drug response, whereas mechanical sex-differences are also underpinned by anatomical differences.

**Main Contributions:**

- Development, calibration and validation of sex-specific human ventricular electromechanical, multiscale models.
- An analysis of clinical randomised trial data in the context of sex-specific effects of multi-channel blockers on the ECG by dosage, where previous analysis focused on pharmacokinetics.
- Consideration of both sex-specific electrophysiology and anatomy explains sex-specific differences on the impact of drugs on ECG and mechanics.
- Simulations demonstrate that the reduced repolarisation reserve in females increases the susceptibility to QTc prolongation via potassium channel block compared to males, and more robust calcium dynamics protect against t-wave amplitude reduction and the more severe contractility loss through L-type calcium inhibition observed in males.

## Introduction

Age-adjusted mortality for cardiovascular diseases has declined less in women than in men, and remains a leading cause of mortality [1]. A critical factor in this difference is the under-representation of women in basic research and clinical studies [2], extending to new approach methodologies (NAMs) such as human-based computational modelling and simulation [3]. This neglect overlooks crucial differences in electrophysiology, anatomy and mechanical function, affecting disease manifestation, progression and treatment efficacy [4]. Recognizing sex as a biological variable is essential for personalized treatments. Unravelling the key factors underlying sex-differences is required to improve risk stratification and treatment outcomes for patients and assist health care professionals [1].

Both preclinical and clinical methods are used in the evaluation of medical therapies. Preclinically, a move towards human-based NAMs using *in vitro* and *in silico* approaches is opening up possibilities for consideration of sex-differences in human pathophysiology in mechanistic studies [5, 6]. One of the most advanced areas is cardiology, with human heart models for electrophysiology [7], mechanics [8, 9], electromechanics [10] and haemodynamic function [11]. A large body of research has established their credibility through agreement with experimental and clinical data from subcellular processes to clinical biomarkers from ECG and imaging [12]. These advancements allow for virtual testing of therapies using modelling and simulation, through in silico clinical trials and digital twins, as it is standard in other industries such as the automobile industry [13]. More recently, inclusion of sex-differences in human-based electrophysiology models has been demonstrated [14, 15], but existing studies focus on cell and tissue simulations or electrophysiological-only simulations which excludes the possibility for inotropic investigations or mechanistic analysis of sex-differences in drug response and disease from ionic changes to clinical endpoints.

To this end, we developed, calibrated and validated sex-specific human ventricular electromechanical multiscale models with the goal of enabling mechanistic investigations of sex-differences in therapy response from subcellular to whole-organ dynamics. As a novelty, both electrophysiological and mechanical function of the human heart is considered in our study, to enable joint investigations into standard clinical biomarkers such as ECG and ejection fraction. We demonstrate that integrating experimental and clinical data as well as biophysical knowledge into the models allows predicting and explaining sex-differences in treatment responses [16]. Furthermore, mechanistic investigations unravelled the key subcellular process determining sex differences in drug response. Such investigations could augment preclinical studies and clinical trials by providing a mechanistic, sex-specific testbed for therapy evaluation.

## Methods

### Sex-specific cell models of human ventricular electromechanics

The human ventricular cellular electromechanical model, ToR-ORd-Land, described in Figure 1 [7, 8] was used, due to its established credibility in cellular and biventricular electromechanical simulations [9] [17] and its credible use for *in silico* trials [18]. Its simulated output includes the action potential, calcium transient and active tension, as well as all ionic currents and excitation-contraction coupling dynamics.

**Figure 1 –.**
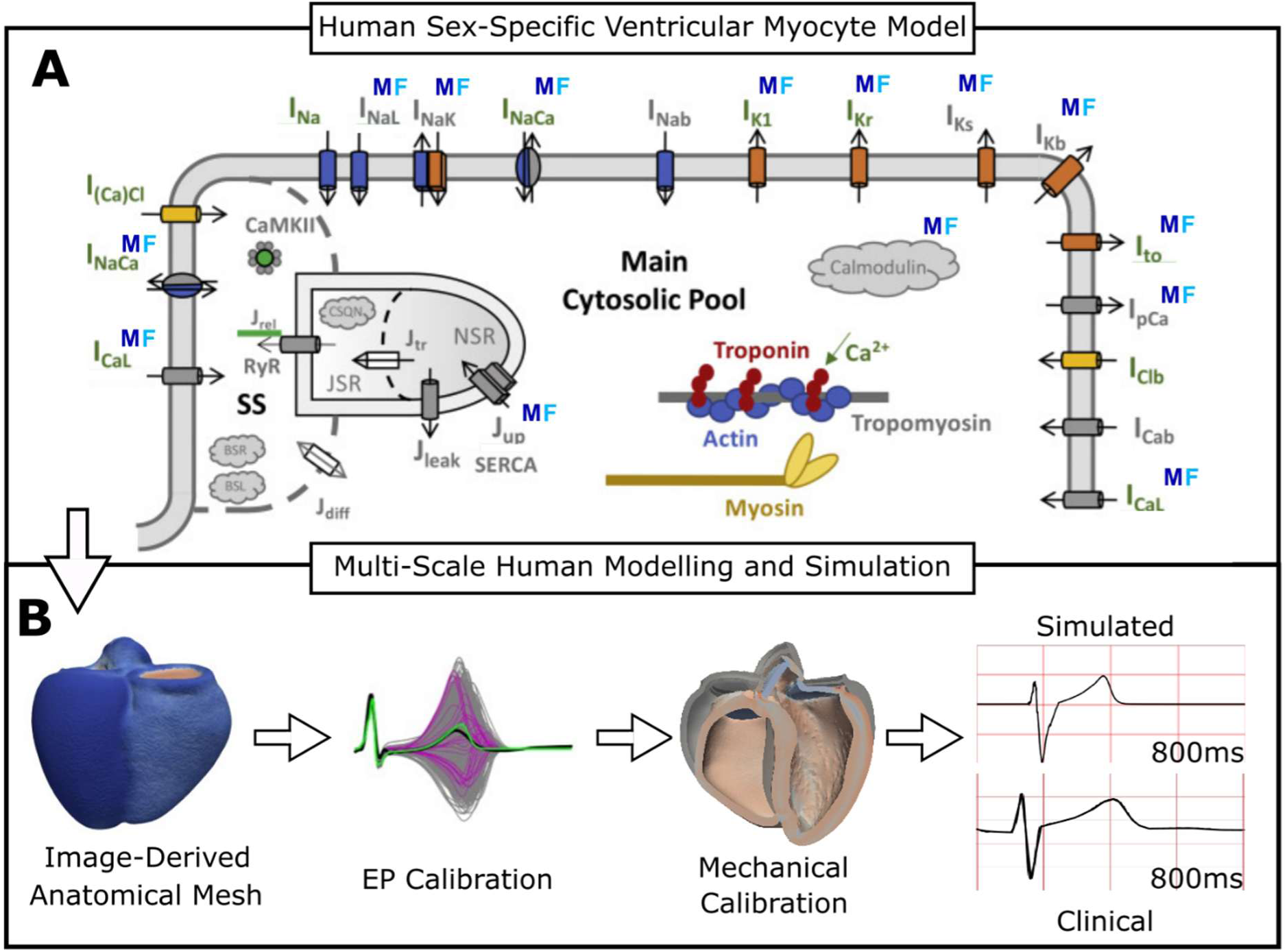
Multi-scale sex-specific modelling and simulation framework. **A:** Schematic representation of ionic conductance, Ca^2+^ dynamics, buffering and active tension dynamics in the human electromechanical ventricular cell model. Variables which have been adjusted for sex-specific differences are labelled by “M” and “F”. **B:** Multi-scale simulation framework incorporating sex-specific anatomy and electromechanics from A, electrophysiology (EP) as well as mechanical and ECG-based calibration.

To calibrate for sex-specific behaviours, ionic conductances were adjusted using relative mRNA expression ratios of ion channel markers from non-diseased adult human male and female ventricular myocytes (Supplementary Material, SM1) [19]. This method aligns with previous electrophysiology-only simulation studies [14, 15]. Activity ratios for these channels (Figure 1; Table 1) were calculated using this data, compensating for gender bias present within the model, estimated to be approximately 60% male based on a linear interpolation of the gender ratio of human donors (N = 140; 78 M) [20] and the sex of additional data. Initially, adjustments were made to the male endocardial model, followed by derivations of mid-myocardial, epicardial and female variants through conductance scaling factors. These adjustments primarily focused on balancing inward potassium channels, calcium handling and calmodulin buffering.

**Table 1 –.**
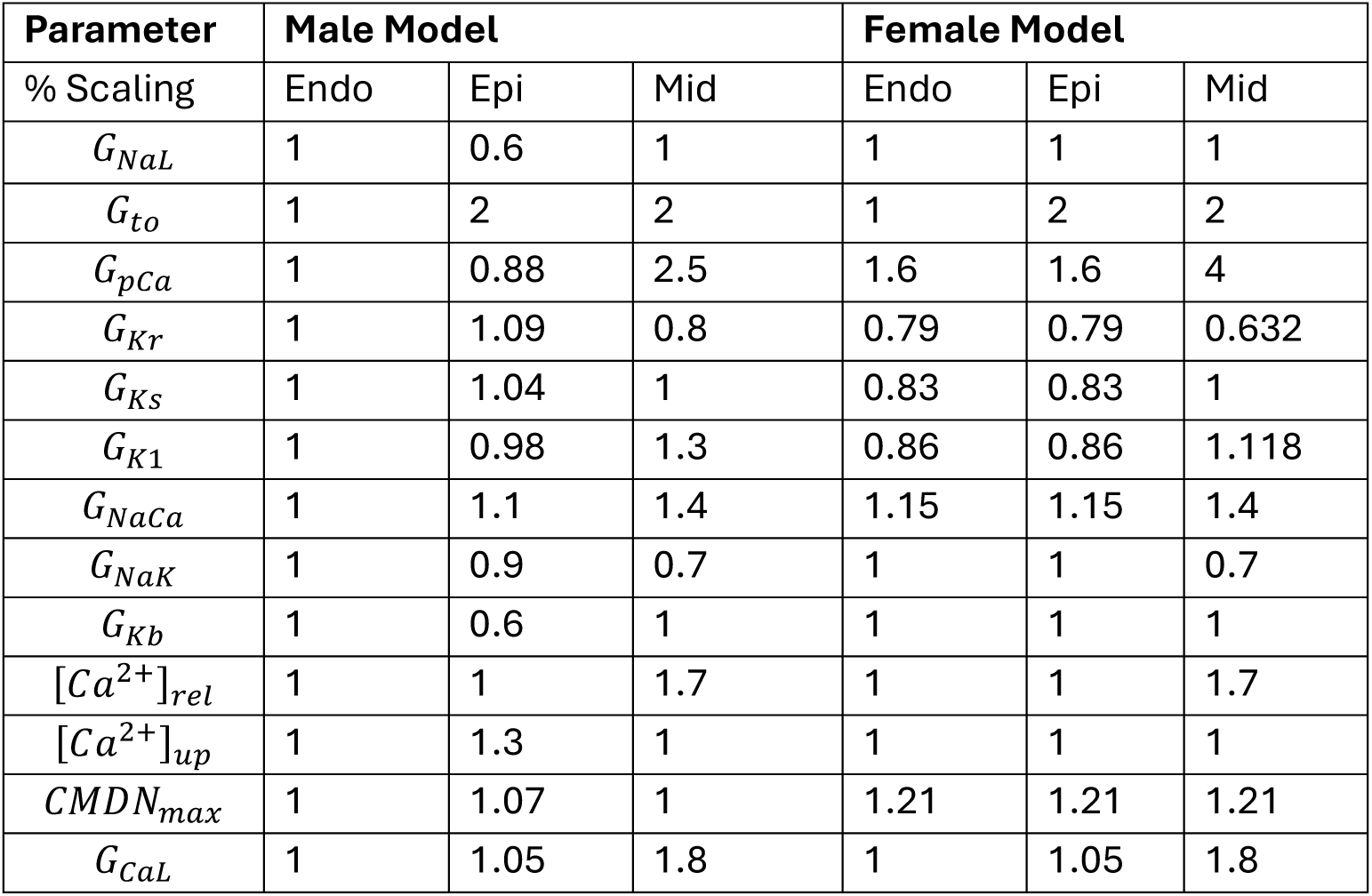
Sex-specific subcellular ion current activity ratios. Sex-specific cell type scaling from the recalibrated baseline male endocardial model, derived from mRNA and protein ion subunit expression data in SM Table 1 [19, 21, 22], previous ORd scaling modifications [14, 15] and existing sex biases.

| Parameter | Male Model |  |  | Female Model |  |  |
| --- | --- | --- | --- | --- | --- | --- |
| % Scaling | Endo | Epi | Mid | Endo | Epi | Mid |
| $G_{NaL}$ | 1 | 0.6 | 1 | 1 | 1 | 1 |
| $G_{to}$ | 1 | 2 | 2 | 1 | 2 | 2 |
| $G_{pCa}$ | 1 | 0.88 | 2.5 | 1.6 | 1.6 | 4 |
| $G_{Kr}$ | 1 | 1.09 | 0.8 | 0.79 | 0.79 | 0.632 |
| $G_{Ks}$ | 1 | 1.04 | 1 | 0.83 | 0.83 | 1 |
| $G_{K1}$ | 1 | 0.98 | 1.3 | 0.86 | 0.86 | 1.118 |
| $G_{NaCa}$ | 1 | 1.1 | 1.4 | 1.15 | 1.15 | 1.4 |
| $G_{NaK}$ | 1 | 0.9 | 0.7 | 1 | 1 | 0.7 |
| $G_{Kb}$ | 1 | 0.6 | 1 | 1 | 1 | 1 |
| $[Ca^{2+}]_{rel}$ | 1 | 1 | 1.7 | 1 | 1 | 1.7 |
| $[Ca^{2+}]_{up}$ | 1 | 1.3 | 1 | 1 | 1 | 1 |
| $CMDN_{max}$ | 1 | 1.07 | 1 | 1.21 | 1.21 | 1.21 |
| $G_{CaL}$ | 1 | 1.05 | 1.8 | 1 | 1.05 | 1.8 |

### Human multiscale modelling and simulation framework

The human multiscale modelling and simulation framework is described here in brief; adapted from the electromechanical model described in Wang et al. (2021) [9]. Two biventricular anatomical meshes, female and male, (Figure 1B) were generated from CT images taken during the R-R interval in healthy subjects by Rodero et al. (2021) [23]. Simulations of electrical propagation in each biventricular anatomy were conducted using the monodomain equation. Orthotropic diffusion along the fibre, sheet and sheet-normal directions was implemented [24]. Ionic current and intracellular Ca^2+^ dynamics were modelled using the sex-specific cell models described above. Sinus rhythm was simulated through stimulation of a fast-activation endocardial layer, representing Purkinje-myocardial junctions with root node locations to achieve realistic QRS morphology. [25]. Standard 12-lead electrode positions were mapped to both anatomies from the geometry of a previous study [26], and the ECG was simulated using the pseudo-ECG method from these node positions to enable comparisons. Apex-to-base *G_Ks_* scaling was calibrated using a fast-inference method [27] (supplementary material SM2), to achieve physiological QRS and T-wave morphology with the non-specific ionic model using the same healthy ECG on for both anatomies, to allow for identifying the contribution of ionic-level sex-differences and drug effects. [26].

Contractile force generation was simulated using the human-based active tension Land et al. model [28], coupled to the sex-specific ionic models, as in [8]. Strongly-coupled electromechanics were modelled as in [10], with orthotropic passive mechanical behaviour and linear momentum balanced with inertial effects. Furthermore, a five-phased state machine was added, with an active diastolic inflation phase following isovolumetric relaxation, in which elastic recoil may occur. A description of mechanical boundary conditions can be found in supplementary material SM3, based on [9], and a detailed mathematical description of the cardiac baseline mechanical model is available in [10, 11]. Parameters for passive mechanics were mostly adapted from [9]. Peak active tension, pericardial stiffness and the resistance of the arterial Windkessel model were calibrated to achieve physiological ejection fractions matching both healthy adult males and females in the UK Biobank (n= 806, 46% Male) [29]. The same linear parameters of passive mechanical behaviour and the Windkessel model were used for both sexes. A list of all parameters can be found in supplementary material SM3. Electromechanics were simulated for five beats of 1000ms cycle length, achieving ECG and pressure-volume convergence after the second beat (supplementary material SM4).

### Calibration and validation strategies

Verification, validation, and uncertainty quantification (VVUQ) are critical for ensuring the credibility of simulation outputs. We follow the example set forth for the ASME VCV40 guidelines applied to the context of cardiovascular science [30].

#### Question of interest

How accurately human multi-scale, electromechanical simulations reproduce and explain sex-specific differences in cellular, ECG and pressure-volume loop biomarkers in control and under pharmacological intervention of Dofetilide and Verapamil?

#### Context of use

Multi-scale electromechanical human ventricular simulations allow for mechanistic investigations to evaluate (patho)physiological therapeutic interactions. We focus on evaluating the model in healthy conditions to establish credibility for sex-specific studies.

#### Calibration and validation strategy for the single cell model

At the single cell level, calibration was performed by adjusting ionic conductance scaling to match sex-specific protein genomic expression data. Minor adjustments to Ca^2+^ handling parameters were performed as described prior, to match Ca^2+^ characteristics in each sex-model to sex-specific experimental data describing biomarker ranges for action potential. Following this, both models were benchmarked against the baseline unisex/generic ToR-ORd-Land electromechanical cell model and compared with experimental reference ranges for adult human action potential (AP), calcium transient and active tension biomarkers [8].

Validation was performed through reproduction of drug modulation in AP, calcium transient and active tension via the multi-channel blockers Dofetilide, Ranolazine, Quinidine and Verapamil. Simulated outputs were compared with independent experimental data.

In addition, a population of models (n = 1000) was generated by varying ionic conductance scaling in the ranges 50-150%, independent of sex-based scaling factors, filtered using healthy biomarker ranges for model outputs to simulate the range of physiological variability in males and females [31].

#### Calibration and validation strategy for human biventricular models & simulations

Sex-specific biventricular models were calibrated, building on [9], to achieve peak active tension, pericardial stiffness and arterial resistance and sex-stratified clinical ECG (QRS, QT, T-wave) and pressure-volume (ejection fraction (EF), end-diastolic volume (EDV), stroke volume (SV) biomarkers consistent with data for a healthy cohort of adult patients from the UK Biobank [29, 32]

Validation of biventricular models and simulations was conducted through comparison to clinical ECGs showing pharmacological modulation of QTc and T-wave morphology, comparing dose-dependent electrogenic and mechanical response compared with independent randomised controlled trial data and experimental data.

Clinical data used for comparison were obtained from ClinicalTrials.gov ID NCT01873950, a randomised clinical trial involving 22 healthy adults (11 male, 11 female) aged 18-35 with a BMI of 18-27 kg/m^2^ [33]. A follow-up study analysed sex differences in the pharmacokinetic profiles of Verapamil, and the pharmacodynamic action of Dofetilide on the QTc interval and T-wave amplitude [34]. ECG recordings obtained from this dataset were re-analysed to investigate dose-dependent sex differences by calculating molar plasma concentrations of each compound at each timepoint. Changes in QTc interval and T-wave amplitude were determined by matching patient timepoints with their placebo measurements and calculating the difference of the means for three recordings between each timepoint with an associated plasma concentration and the placebo.

The sex-specific ionic models and four randomly selected subjects from cellular populations of models were used for biventricular simulations to assess variability in ECG and mechanical parameters. We quantify uncertainty through a sensitivity analysis of key potassium, calcium and sodium current conductance on the QT duration and T-wave (supplementary material SM7). The impact of ECG personalisation on ECG biomarkers is also detailed in (supplementary material SM8). In addition, a sensitivity analysis of passive (non-anatomical) mechanical parameters (Supplementary Materials, SM9) was performed to assess any impacts of mechanical differences, whether through sex-specific or pharmaceutical modulation, would impact the ECG (supplementary material SM9).

### Simulation of Pharmacological Action

Drug effects were modelled by blocking the relevant ionic currents in the cell model, as in patch-clamp experiments [35–37] using a simple pore block model (supplementary material SM5). Biventricular simulations were performed with Dofetilide (1 nM to 7 nM), and Verapamil (0.05μM to 0.35μM) corresponding to the range obtained in pharmacokinetic profiles of patients in the trial [33].

### Simulations

Cellular simulations were performed using MATLAB (Mathworks Inc. Natwick, MA, USA) with the ODE solver *ode15s*. Steady state was achieved at 1Hz pacing before computing biomarkers. Drugs were virtually applied after reaching steady state, and simulations ran for another 200 beats at the specified pacing frequency. The final 10 beats were analysed for repolarisation abnormalities and after-contractions, with all biomarkers taken from the last beat in normal sinus rhythm.

Biventricular simulations were conducted using the high-performance numerical software, Alya, for complex coupled multi-physics, multi-scale problems [11] on the supercomputer ARCHER2, the UK National Supercomputing Service (UKRI, EPCC, HPE, Cray and the University of Edinburgh). Biventricular simulations were performed on 128 CPUs with a total of 1536 cores, for 4 hours to simulate 3 beats. The electrics and mechanics were solved implicitly as different modules to achieve global convergence every timestep.

### Data Availability

The simulation input files and Alya executables required to replicate the simulated results presented in this manuscript will be made available upon request. Python and MATLAB scripts for the cellular models, simulations and post-processing of the biventricular simulations can be found at: https://github.com/MaxxHolmes/Sex_Specific_Human_Electromechanics. Additionally, code used to delineate the simulated ECGs can be found at: https://github.com/MaxxHolmes/ECG_Delineation.

## Results

### Calibration of Sex-Specific Human Cellular Ventricular Models

Simulations using sex-specific cardiomyocyte models recapitulate the prolonged APD observed in females vs. males (17% longer in endocardial cells) consistent with clinical QTc recordings of healthy adults (Figure 2A) [38], and human failing myocytes [39]. Simulations also reproduced lower calcium transient amplitude in females (13% smaller in endocardial) observed in patients [40], and more rapid calcium transient decay in females (7% longer in endocardial), observed in murine experiments and patients [40, 41]. No marked sex differences were observed in the simulated diastolic calcium levels or SR calcium content, aligning with measurements in patients [40]. Primary ionic differences are summarised in Figure 2C; inward potassium rectifier currents are generally reduced in the female model, whereas *I_NaCa_* and *I_pCa_* are enhanced. Parameter values are presented in Table S2.

**Figure 2 –.**
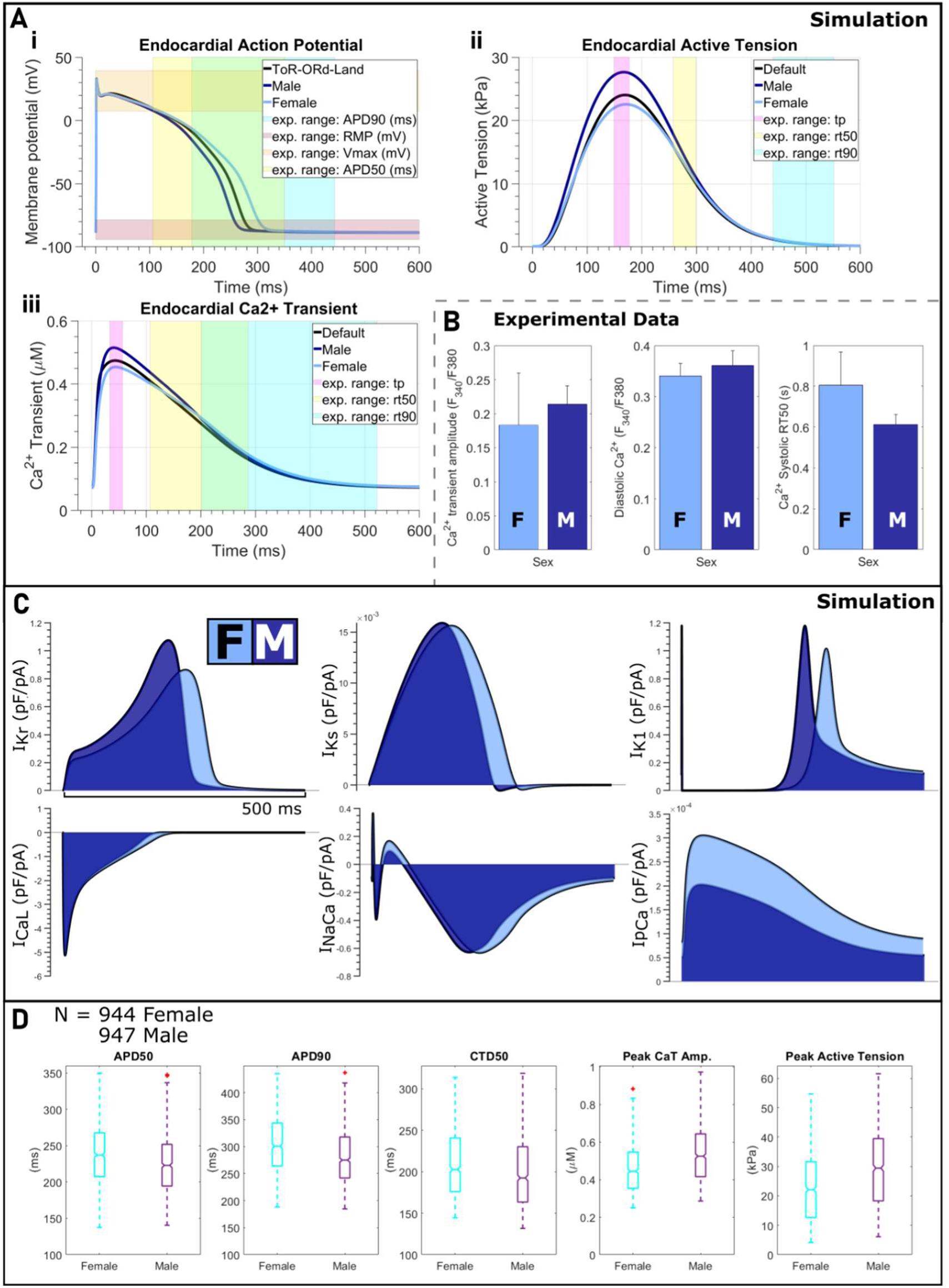
Human female and male cellular electromechanical models calibration: **A:** Simulated **i)** action potential (AP), **ii)** active tension and **iii)** intracellular calcium transient traces pacing at 1Hz for the baseline non-specific model (black), the female model (cyan) and the male model (navy). Experimentally obtained ranges for key biomarkers are represented as transparent blocks as denoted in the legend. **B:** Experimental data (right) showing sex-differences in calcium transient biomarkers which match our simulated outputs [40]. **C:** Ionic current traces for female and male cell models overlaid. **D:** Biomarkers considering variability in population of female and male models, with ionic conductance scaled between ± 50% of sex-specific values in panel C.

Figure 2D further demonstrates that, in females-specific simulations, prolonged APD and calcium transient, as well as larger calcium transient amplitude and active tension, are consistently present in populations of female (N = 944) and male (N = 947) models incorporating variability in ionic currents.

### Validation of Sex-Specific Cellular Models by Drug Action

Simulations with sex-specific cellular models were conducted with four compounds: a specific hERG blocker, Dofetilide; a primary calcium blocker and secondary hERG blocker, Verapamil; a primary sodium and secondary hERG blocker Ranolazine; and Quinidine, a multichannel calcium, hERG and sodium blocker (Figure 3), as described in Supplementary Material (SM5).

**Figure 3 –.**
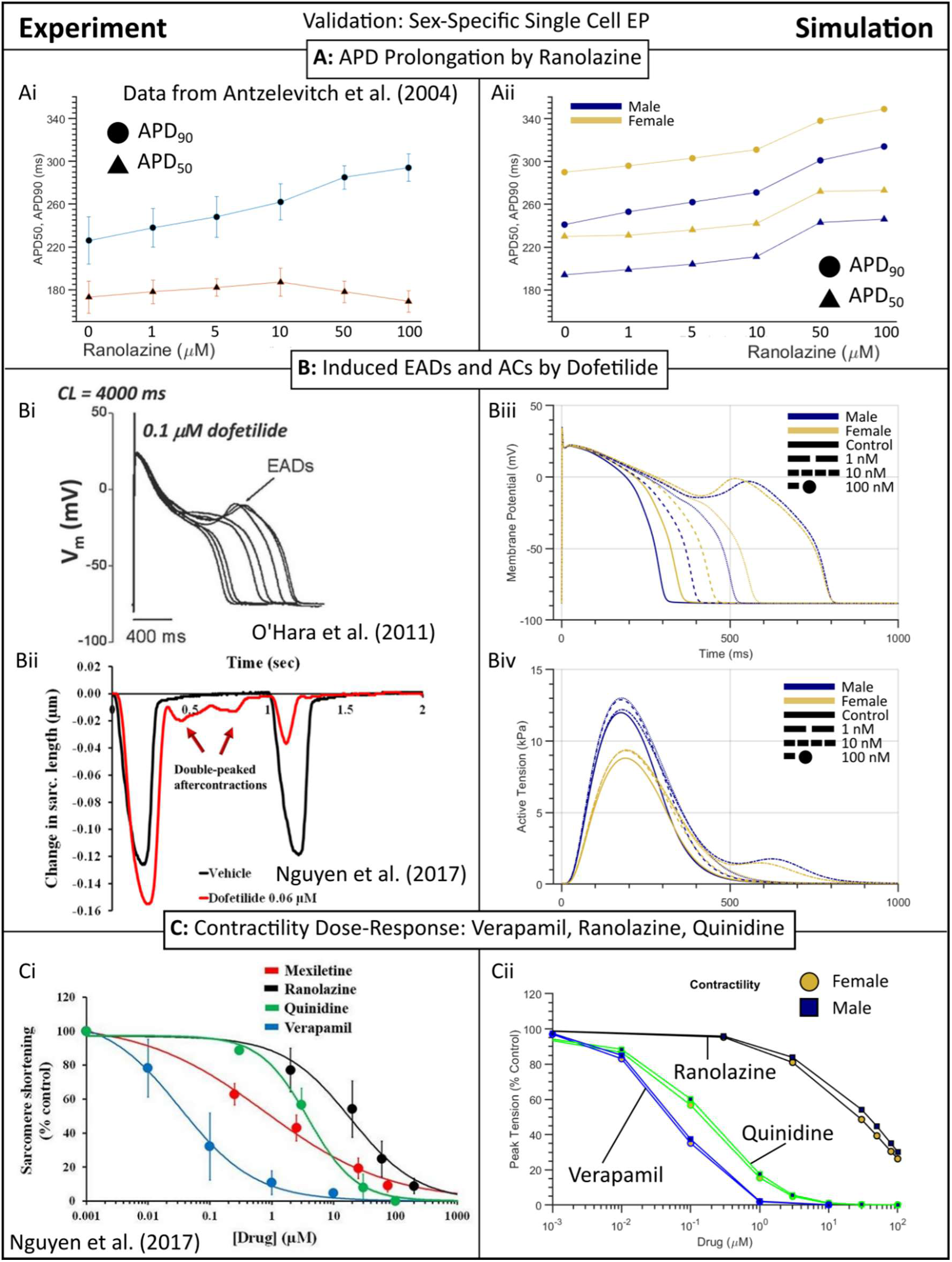
Sex-Specific Validation of drug-induced effects on Action Potential and Contractility. **A:** APD prolongation by Ranolazine. **i**) APD_90_ and APD_50_ (ms) data from canine LV wedge preparations *[3C]*. **ii)** Simulated APD_90_ and APD_50_ for male and female sex-specific cell models. **B:** Induced early-after depolarisations (EADs) and aftercontractions (ACs) by Dofetilide. **i)** AP traces and EADs in undiseased human tissue in control and under 100nM of Dofetilide [20]. **ii)** Sarcomere shortening in adult human ventricular myocytes in control and after exposure to Dofetilide [42]. **iii)** Simulated AP traces and **iv)** active tension traces in control and under 1, 10 and 100nM of Dofetilide. **C:** Contractility dose-response. **i)** Sarcomere shortening recorded in human myocytes for Ranolazine, Quinidine and Verapamil [42]. Mexiletine also pictured. **ii)** Simulated peak tension by drug concentration.

The single-cell simulations replicated the action potential duration (APD) prolongation induced by Ranolazine and Dofetilide observed in human ventricular myocytes [20] and in canine ventricular wedge preparations [36] (Figure 3A, B). The female cellular model showed greater APD_90_ prolongation under both compounds, leading to early-after depolarisation (EAD) initiation, due to reduced repolarisation reserve compared to the male model. Simulated APD_50_ followed the same trend up to 10 µM but continued to increase whereas the experimental data decreased at higher doses. At Dofetilide concentrations that evoked EADs in human cells, the male and female model both evoked EADs and after-contractions [20, 42] (Figure 3B). The simulated dose-dependent contractility responses for Verapamil, Ranolazine, and Quinidine agreed with observed contractility profiles in human ventricular myocytes, with minor differences between the male and female models explained by the larger baseline active tension generation and intracellular calcium concentrations in males (Figure 3C).

### Sex-Specific Biventricular Simulations at Baseline

Figure 4 illustrates the mechanical and ECG outputs of the sex-specific simulations compared with healthy population values obtained from the UK Biobank [32]. QRS durations were calibrated between 80 and 100ms (Simulated Female: 80ms, Male: 95ms in all models) to be consistent with healthy QRS durations from the same population. Mean T-wave amplitudes for each sex-specific simulation were positive across all pre-cordial leads for both male and female simulations.

**Figure 4.**
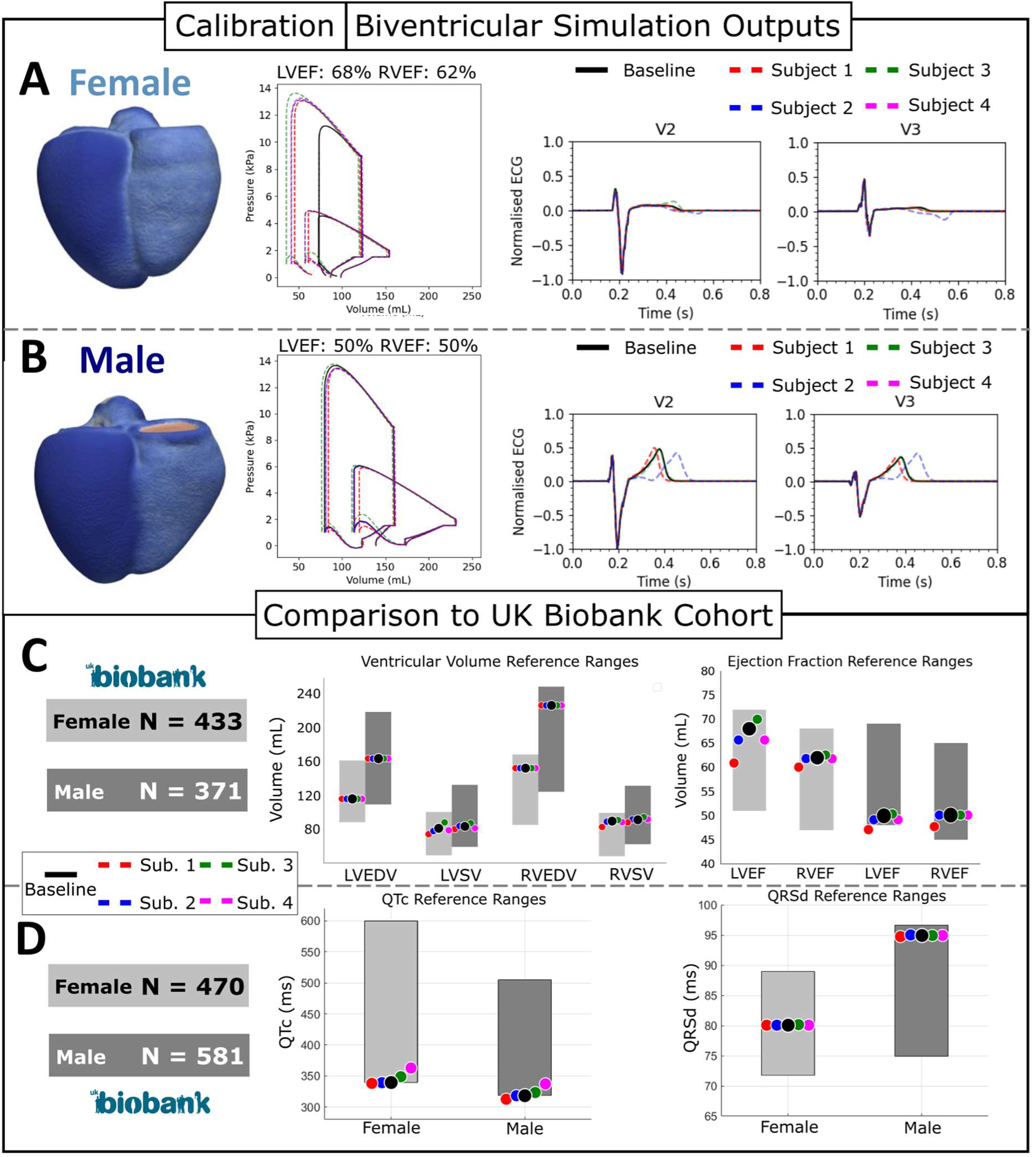
Simulated Clinical Biomarkers with Sex-Specific Models and Comparison to UK Biobank Cohorts. **A**: Female and **B**: male models with respective sex-specific electrophysiology and anatomies reproduce physiological mechanical and ECG outputs at a pacing rate of 75 Hz. **C**: Simulated ventricular end-diastolic volume (EDV), stroke volume (SV) and ejection fractions (EF) in the left (LV) and right ventricle (LV) compared with sex-stratified reference ranges from a healthy adult UK biobank cohort *[2S]*. **D**: Simulated QTc (ms) and QRS duration (ms) averaged over precordial leads compared with sex-stratified reference ranges from a healthy adult UK Biobank cohort [32].

In the female simulation (Figure 4A), baseline LVEF and RVEF are 68% and 62% respectively (Across all female models: LVEF: 62% - 70%; RVEF: 60-62%). These values correspond to the upper quartile for healthy adult Caucasian women (Figure 4C). Conversely, the male simulation produces a baseline LVEF and RVEF of 50% in both cavities (Across all male models: LVEF: 47-51%; RVEF: 48-51%) (Figure 4B, C), in the lower quartile for healthy Caucasian men. The higher EF in the female simulation is primarily due to lower end-diastolic volume (EDV) resulting from a smaller heart size and a similar stroke volume (SV). Notably, the female anatomy used in these simulations has a LV myocardial volume 66% smaller than the male anatomy.

Figure 4D shows mean QTc interval (via Fridericia’s correction) for the simulated female (baseline: 340 ms; range: 338 – 366ms) and male (baseline: 312ms; range: 308 – 334ms) are within UK Biobank range, calculated by Smith et al. (2024, [32]) with QT dispersion of 11ms and 9ms respectively across precordial leads.

### Sex-Specific ECG Characteristics explained by Specific Ionic Current Differences

To evaluate the relative importance of electrophysiological versus anatomical sex-differences, we conducted simulations combining both sex-specific electrophysiology in both female and male anatomies (Figure 5).

**Figure 5 –.**
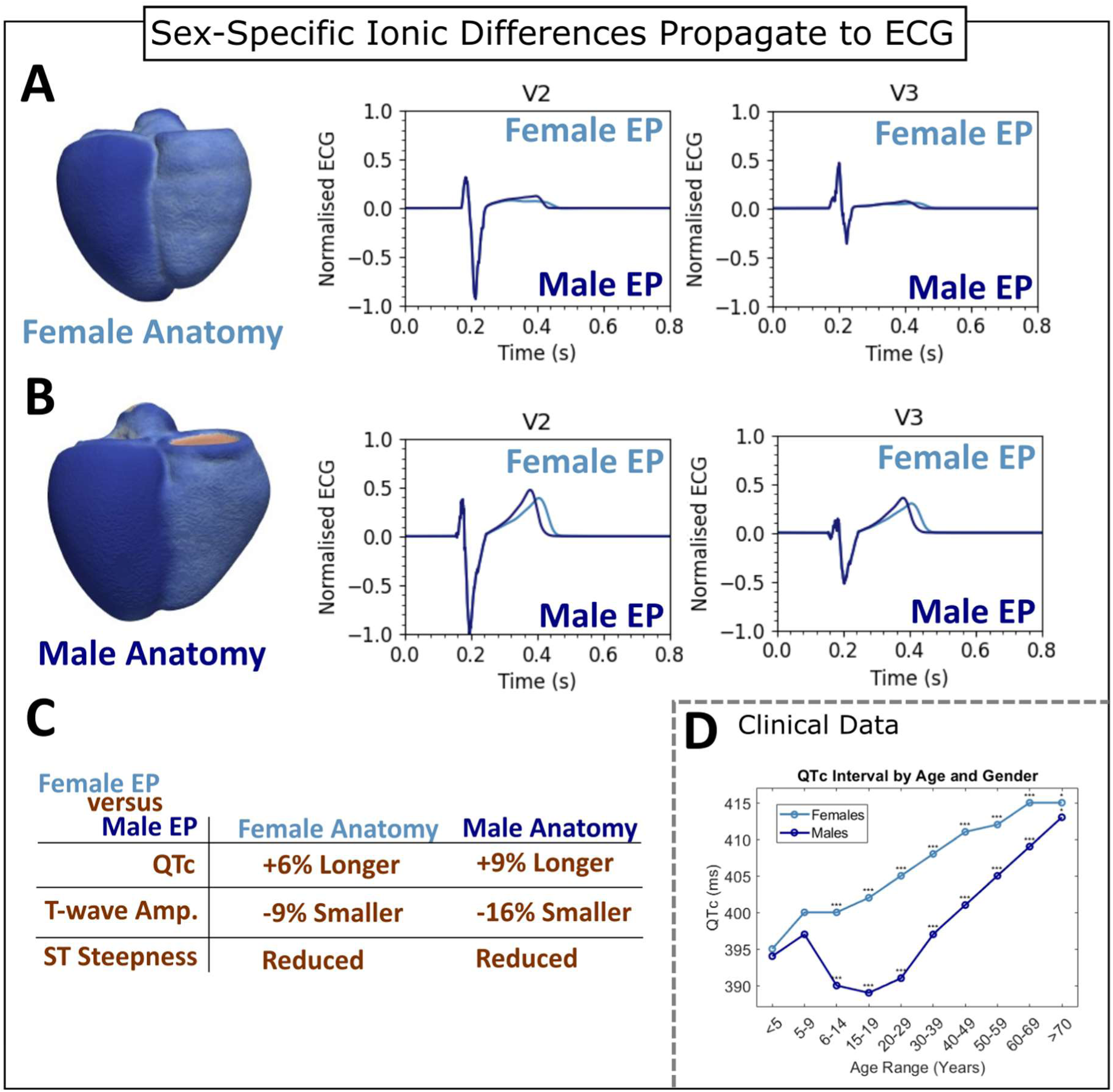
Sex-Specific ECG characteristics: electrophysiology versus anatomy. Simulated ECG on V2 and V3 leads with female (cyan) and male (navy) electrophysiology on **A:** female and **B:** male anatomies. **C**: Measured differences between using female and male electrophysiology on the female and male anatomies. The female electrophysiology is shown to produce a prolonged QTc, smaller T-wave amplitude and reduced ST steepness in both anatomies. **D**: Clinical QTc measurements for healthy patients stratified by age and gender [44].

In the female anatomy (Figure 5A), QTc interval is 6% longer when simulated with female electrophysiology than with male electrophysiology, which can be attributed to the collective action of smaller repolarising K^+^ currents in females. Additionally, simulation with female electrophysiology resulted in a 9% decrease in T-wave amplitude and a shallower ST segment compared to using male electrophysiology. This reduction in T-wave amplitude is due to the less pronounced electrical activity caused by reduced repolarisation reserve in females.

Similarly, using the female electrophysiology with the male anatomy yielded a 9% increase in QTc duration, and a 16.3% decrease in mean T-wave amplitude. These patterns correspond with clinical observations [32, 43]; females typically present a longer QTc at all ages (Figure 5D), and shallower, broader T-waves. A sensitivity analysis of passive (non-anatomical) mechanical parameters (Supplementary Materials, SM9) did not impact the ECG when modulated from 50% to 200% scaling. This suggests that electrophysiological differences between males and females play the dominant role in determining ECG sex-differences.

No pronounced mechanical differences were observed when simulating either anatomy with different sex-specific electrophysiology. As both anatomies were simulated with the same passive mechanical parameters, this suggests that sex-differences in mechanical biomarkers are a result of anatomical differences.

### Sex-Differences in Dofetilide Response

Clinical ECGs (Figure 6A) show QT interval prolongation under Dofetilide, with female patients experiencing significantly larger QTc increases (P < 0.05) at higher pharmacokinetic concentrations compared to males. Our simulated ECGs (Figure 6B) recapitulate this QT prolongation, predicting QT interval changes within the population data bounds from the clinical trial [33]. We observed similar trends in QTc dose-response and sex-specific effects (Figure 6Ci, iii). The male simulation predicts an increase of 60ms (1 nM) to 115 ms (7 nM) compared with an increase in the female of 106ms (1 nM) to 178ms (7 nM), indicating a 54-78% difference.

**Figure 6 –.**
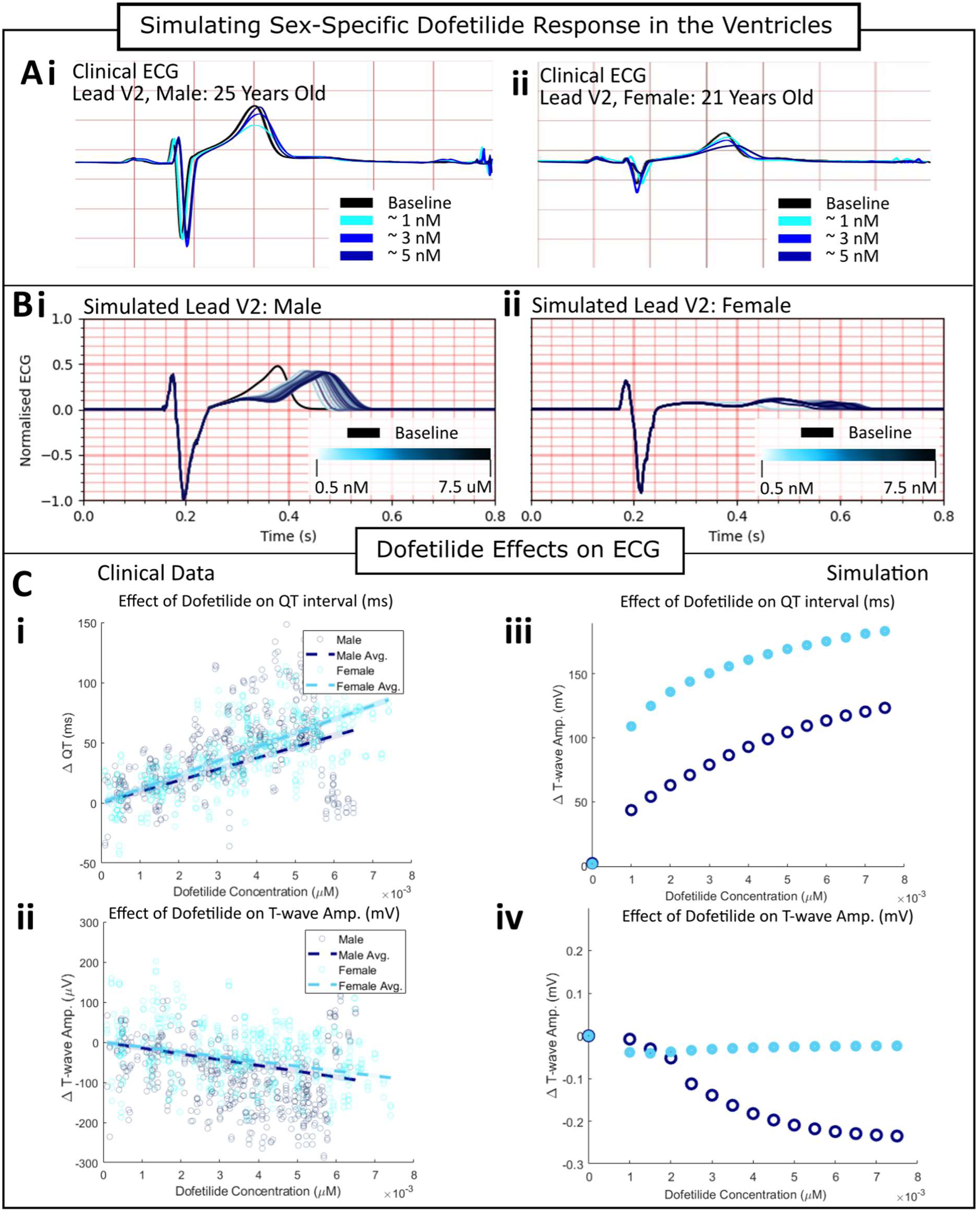
Clinical and Simulated ECG biomarkers with Dofetilide administration. **A**: Clinical ECG recording of V2 lead at baseline and with 1, 3 and 5 nM of Dofetilide for a **i)** 25-year-old male and **ii)** 21-year-old female. **B**: Simulated ECG recording of V2 lead at baseline and under the effects of Dofetilide ranging from 0.5nM to 7.5nM for the virtual **i)** male and **ii)** female. **C:** Clinical ECG analysis of Dofetilide dose-response on **i)** QT interval in ms and **ii)** T-wave amplitude (mV) in the male and female patient cohorts. **D**: Simulated sex-specific ECG dose-response of **iii)** QT interval and **iv)** T-wave amplitude. Values shown are averaged from all precordial leads.

Male and female cohorts in the clinical trial show a dose-dependent decrease in T-wave amplitude when treated with Dofetilide, with males presenting a slightly larger decrease, but not-significant. Simulations produce coherent results with both male and female showing decreases in T-wave amplitude, but only the male simulation shows a dose-dependent reduction in T-wave amplitude. The male simulation shows a reduction from −0.01mV at 1nM, to −0.25mV at 7.5 nM, which is within the range observed in male patients in the clinical trial (Figure 6D). The female simulations however do not show a dose-dependent reduction and has a consistent 0.03-0.04 mV reduction in amplitude across all doses.

This is coherent with our sensitivity analysis (supplementary material, SM7), which suggests hERG channel block leads to a decrease in T-wave amplitude in male electrophysiology with no significant changes in female electrophysiology. It should be noted the female model produces bifid T-waves (observed in the clinical trial in two female subjects with similarly low T-wave amplitudes [34]). The small baseline T-wave in the female model means it is less susceptible to further reductions, and it can be seen in the clinical trial results (Figure 6Cii) that several female patients displayed no change in T-wave amplitude.

The simulation outcomes suggest that sex-differences in response to hERG block are caused by differences in repolarisation reserve. Lower repolarisation reserve through comparatively reduced inward potassium currents in the female electrophysiology makes it more sensitive to further reductions in *I_Kr_* by Dofetilide. The balance of repolarising and depolarising currents shifts towards depolarisation, leading to a higher T-wave amplitude in women. The delayed return to RMP enhances the T-wave’s prominence, extending the QT interval further than in males.

### Validation: Verapamil Response

Simulated ECGs (Figure 7A) showed a minor increase in T-wave amplitude in female simulations, and a decrease in male simulations. Unlike Dofetilide, Vicente et al. [34] did not observe sex-specific trends in Verapamil’s effect, but noted significant differences in pharmacokinetic profile, with female patients having significantly higher plasma drug concentrations at the same dosage administration than male patients.

**Figure 7 –.**
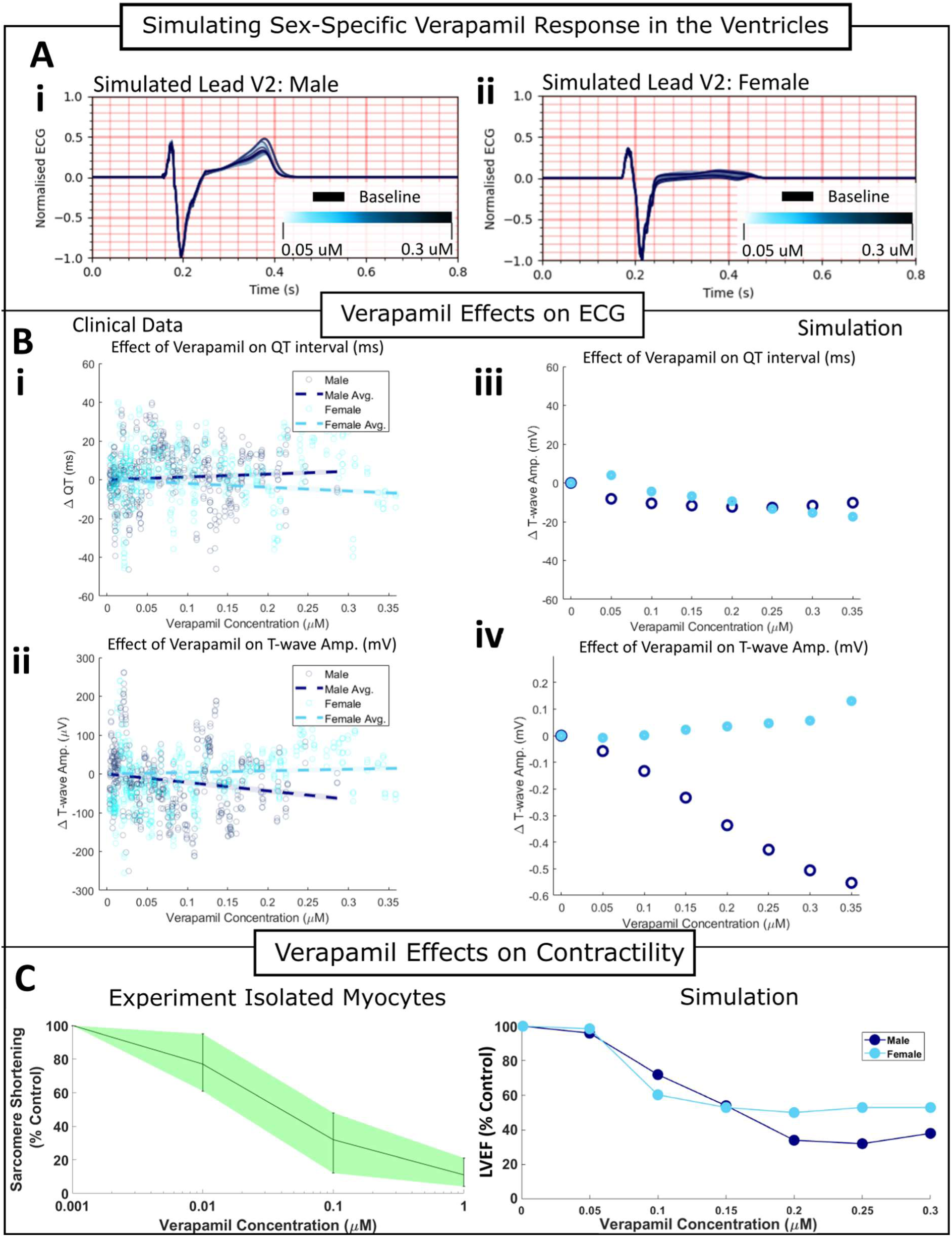
Comparison of Verapamil Effects: Clinical and Simulation. **A:** Simulated ECG recording of V2 lead at baseline and under the effects of Verapamil ranging from 0.05μM to 0.3μM for the virtual **i)** male and **ii)** female. **B:** Clinical data on the effects of verapamil on the male and female cohorts on **i)** QT interval and **ii)** T-wave amplitude. Simulated ECG dose-response of **iii)** QT interval and **iv)** T-wave amplitude for virtual male and female. Values shown are average and dispersion from all precordial leads. **C:** Comparison of experimental data (sarcomere shortening, left) and simulated contractility effects (LVEF, right) under the influence of Verapamil from 0.05μM to 0.4μM.

Clinical data show no significant QTc prolongation in males (Figure 7Bi) and a minor decrease in females, which is mirrored in our simulations (Figure 7Ci). T-wave amplitude changes show divergent behaviour: females in the clinical trial observed no significant change, while males saw a notable dose-dependent reduction under Verapamil. This behaviour can also be shown in the simulated ECG biomarkers; the female simulations show negligible change with increasing dosage, whereas the male simulations demonstrate a dose-dependent decrease (Figure 7Cii).

Contractility effects were compared with experimental data of sarcomere shortening in human ventricular myocytes [42] (Figure 7C). Simulation outputs were sensitive to Verapamil’s effects, showing large losses of contractility and LVEF reduction. The female simulation suggests greater resistance to the effects of Verapamil on LVEF at higher concentrations (Figure 7C).

Verapamil primarily blocks *I_CaL_*, and secondarily blocks *I_Kr_*. The clinical data and simulations both suggest that there is no significant difference to the QT interval as the *I_Kr_*-induced prolongation is balanced by the *I_CaL_*-induced shortening. The T-wave amplitude in males is decreased due to reduced calcium influx, agreeing with previous studies [44], and a stronger reliance on *I_CaL_* for contraction than the female model. The female model has a comparatively enhanced sarcolemma calcium pump, calmodulin and NCX current which supports most robust calcium handling and contractility, which in turn allows for the maintenance of better function under calcium block [45].

## Discussion

### Summary

In this study, we present a sex-specific investigation into human electromechanical cell and biventricular electromechanics from ionic to clinical biomarkers, and their response to pharmaceutical compounds. Credibility is evaluated following established guidelines and specifically through the evaluation of simulation results with experimental and clinical data at cellular and whole-organ level, including successfully reproducing ECG, cellular electrophysiology, arrhythmic mechanisms and contractile profiles at the cellular and organ-scale. Furthermore, we demonstrate agreement with clinical trial data showing sex-differences in response to Dofetilide and Verapamil. The sex-specific models described here represent a mechanistic testbed for virtual evaluation of therapies in females versus males.

We demonstrate that sex-differences in ionic electrophysiology propagate and play an important role in the emergent sex-specific ECG characteristics observed in healthy populations regardless of anatomy and mechanical differences. Specifically, we identify reduced repolarisation reserve in females increasing the susceptibility to QTc prolongation via potassium channel block compared to males, and a more robust calcium reservoir protecting T-wave amplitude reduction and the more severe contractility loss through L-type calcium inhibition observed in males.

### Sex-Specific Models

Women have historically been under-represented in cardiovascular research [2, 3]. Considering known electrophysiological and anatomical sex-differences is essential to understand mechanisms underlying differences in drug efficacy and risk assessment. Previous works on sex-specific human ventricular cell models have utilised electrophysiology-only cell models. This work takes a step forward and unites cell-level phenomena with organ-scale behaviours to investigate how electrophysiological changes at the myocyte scale propagate through to influence organ scale electromechanical behaviours. This work provides a platform to begin to investigate how the mechanisms of age and gender influence the efficacy and safety of pharmaceutical treatments.

Sex differences in ionic currents and cellular electrophysiology are well documented. Females have elongated APD due to lower K^+^ expression and differences in sarcolemmal calcium channel density, exacerbated by oestrogens’ inhibition of the hERG channel and their complex effects on L-type calcium channels [46]. This leads to a lower repolarisation reserve, a comparatively reduced calcium transient amplitude and slower calcium transient decay than male cells [40]. Testosterone enhances the activity of inward potassium channels, and reduces L-type calcium flux in males [46].

The sex-specific cellular models presented here replicate these effects after calibration with ion channel expression data; improving over previous models by including contractility and calcium transients, showing quantitative agreement with experimental data on calcium transient amplitude and decay [40, 41, 45]. Single-cell simulations of sex-specific models show consistent quantitative agreement with experimental data on dose-response in APD morphology and contractility for several multi-channel blockers.

Anatomical sex differences also contribute towards progression and initiation of disease and drug efficacy. The male anatomy has a 66% larger myocardial volume than the female, aligning with observations of greater LV mass and LV wall dimensions in males [4]. Anatomies were selected based on their alignment with UK Biobank reference data for healthy individuals [29]. Both anatomies had their ECG calibrated to UK biobank reference ranges and recapitulate sex-differences in the ECG well; females have a longer QTc interval at all ages regardless of correction method [38], a shallower ST segment and a smaller T-wave amplitude [47] which are all reproduced. Ventricular volume and ejection fraction were also calibrated to sex-stratified UK Biobank reference ranges for healthy Caucasian adults [29].

### Drug Response in Biventricular Electromechanical Models

Vicente et al. [34] analysed the clinical trial used as a comparator in this study [33], and present evidence of sex-specific differences in response to Dofetilide, and the pharmacokinetics of Verapamil. Simulations quantitatively reproduced QT interval modulation for both Dofetilide and Verapamil in both sexes, with qualitative matches in T-wave amplitude for Verapamil and Dofetilide, reflecting sex-specific differences from the clinical trial data.

Simulations and sensitivity analysis support that these sex-differences are due to the relative balancing of repolarisation reserve and depolarising currents, and how the two drugs affect this balance. Dofetilide produced greater QTc prolongation in female simulations due to a comparatively lower repolarisation reserve, resulting in a more prominent T-wave and henceforth a longer relaxation. This correlates with clinical evidence that Dofetilide is less safe in women due to a higher risk of torsade de pointes (TdP) [48]. Verapamil strongly blocks L-type calcium, in addition to the hERG current, which resulted in simulated reduction of T-wave amplitude in males but not in females. This is observed to be due to more robust female calcium handling, via enhanced NCX and sarcolemmal calcium compared to the males [45]. This also plays a role in the reduced loss of contractile function in comparison with males.

### Sex-specific models Credibility

Our sex-specific cell models reproduce sex-characteristics observed in ionic current profiles, AP repolarisation, calcium transient duration, amplitude and active tension generation at the cell level. Drug effects were accurately reproduced for four compounds blocking multiple channels, establishing credibility for the sex-specific cell models, following the examples by Tomek et al. (2019) [7] and Margara et al. (2021) [8]. Sex-specific ECG characteristics were reproduced through the inclusion of this sex-specific electrophysiology at the cellular level, which highlights the biophysical accuracy of our models to propagate these differences to the multi-scale. Additionally, we also quantitatively reproduce alterations to QT interval under the action of Dofetilide and Verapamil and demonstrate the ability to mechanistically investigate differences in predicted changes to the ECG. Sensitivity analysis further supports our conclusions, showing that differences in mechanical parameters do not impact our mechanistic explanations.

### Limitations

Calibration of the sex-specific ionic models was constructed using human mRNA expression data in un-diseased hearts, inheriting the challenge of translating this data to ion channel behaviour as discussed by Walmsley et al. [49]. Some minor alterations to calcium handling dynamics (+- 5% maximum scaling for these parameters) were made to ensure the single cell output matched with reference ranges in healthy adults. Additionally, we considered only differences in conductance, and not, for example, differences in gating dynamics.

The biventricular simulations in this study are constructed with sex-specific anatomies and ionic models, however there are no differences in mechanical calibration due to the lack of sufficient data at the time of the study. To address this, we performed a sensitivity analysis that showed negligible effects of mechanical parameters on the ECG (as in Levrero et al. [10]), and thus we can be confident that our ECG outputs would not change with the inclusion of mechanical differences.

The QT durations in our simulated models are within the population data, but at the shorter quartile for both males and females. The ECG calibration process utilised in this study and developed in our group focuses on the QRS interval and producing realistic T-wave progression. This is something we aim to address in future work with a calibration procedure which takes QTc into consideration.

It is known that sex-differences exist in the mechanical function of the heart, including differences in LV/RV global radial strain, long-axis strain, kinetic energy of the blood and cycle-average vorticity [50]. These factors were not taken into consideration and may influence the mechanical outputs of the simulation, highlighting the importance of including these factors in future works. The filling phase of the PV loop is not flat, as observed in clinical PV loops. This can be improved, however does not impact the results of this study.

In this study two anatomical meshes were utilised due to computational tractability at the whole-organ level and the considerable resources needed for development. This was briefly discussed in Methods. The ARCHER2 calculator suggests that the notional cost for this project, including development was approximately £3,840, with 440 kgCO_2_e and a real terms simulation time of 1600 hours. The inclusion of more anatomies, additional subjects from the population of models or additional compounds would provide additional confidence and statistical power in the analysis of these results, at a proportionally increased cost. In future works, this would be a necessary consideration as the field of *in silico* trials develops and becomes more efficient.

## Conclusions

In this study, we present human sex-specific biventricular electromechanical models of electrophysiology and anatomy for simulations of human electromechanical function and the in-silico drug mechanistic investigations of. We demonstrate agreement with experimental, clinical and population data both ECG and ejection fraction for both sexes, and unravel mechanisms underlying sex-differences in treatment response. The sex-specific electromechanical models presented offer a tool for further investigations into sex-specific differences in disease and treatment response.

## Conflicts of Interest

The authors declare that the research was conducted without any commercial or financial relationships that could be construed as a potential conflict of interest.

## Author Contributions

**MH:** Conceptualization, Methodology, Software, Investigation, Formal Analysis, Validation, Visualization, Writing – original draft, Writing – review & editing. **ZJW, RD, JC:** Methodology, Software, Formal Analysis, Writing - review & editing. **HMN, JT:** Methodology, Software, Writing – review & editing. **HS:** Resources, Software, Formal Analysis, Writing – review & editing. **BR:** Conceptualization, Resources, Writing - review & editing, Supervision.

## Acknowledgements

This work was funded by a Wellcome Trust Fellowship in Basic Biomedical Sciences to Blanca Rodriguez (214290/Z/18/Z) and the CompBioMed Centre of Excellence in Computational Biomedicine (European Commission Horizon 2020 research and innovation programme, grant agreement No. 823712. The computation costs were incurred through the CompBioMedX project (e769), which provided access to ARCHER2, a UKRI national supercomputing service.

For the purpose of open access, the author has applied a Creative Commons Attribution (CC BY) public copyright licence to any Author Accepted Manuscript version arising from this submission.

The authors also thank Dr Mariano Vázquez and the other members of ELEM and the Barcelona Supercomputing Centre for providing research access to the Alya simulation software used in this study.

## Notes

### Competing Interest Statement

The authors have declared no competing interest.

