## Supplementary Materials for "Sex-specific human electromechanical multiscale in-silico models for virtual therapy evaluation"

SUPPLEMENTARY MATERIAL

### SM1. Implementation of Sex-Specific Cellular Differences

Differences in channel activity between the male and female ventricular myocytes were implemented as ionic conductance scaling factors calculated using the expression of genes encoding ion-channel and transporter subunits in epicardial and endocardial tissue samples from non-diseased transplant donors, published by Gaborit et al. [1]. In addition, the ratio of male to female data used in the construction of the ToR-ORd-Land model [2] was taken into consideration when adjusting the conductance of ionic currents in the baseline non-specific model. This was taken as a linear interpolation of the gender ratio of the human donors utilised in the construction of the ORd model [3], and the additional data sources which were utilised to further develop the ToR-ORd model [4], and furthermore the Land et al. contractility model [5]. This is estimated to be approximately 60% male and 40% female. A similar approach to producing sex-specific models was taken by Clancy and Yang (2012) [6] and Peirlinck et al. (2021) [7]; which was utilised when configuring the mid-myocardial cell type. A table expressing the differences in gene expression taken into account is presented in Table S1.

| Biomarker | Species | Evidence | Source |
|---|---|---|---|
| $Na_V1.5$ expression (%) | Human | No male-female differences,<br>250% (± 20%) higher in endocardial cells. | [1] |
| $Ca^{2+}$ ATPase 4 expression (%) | Human | +45.2(± 18.1)% in female epicardial cells<br>+44.9(± 20.6)% in female endocardial cells. | [1] |
| hERG expression (%) | Human | -39.6(± 11.3)% in female epicardial cells<br>-31.4(± 12.6)% in female endocardial cells | [1] |
| MinK expression (%) | Human | -34.8(± 11.1)% in female epicardial cells<br>-34.7(± 10.5)% in female endocardial cells | [1] |
| Kir2.3 expression (%) | Human | -50.3(± 8.4)% in female epicardial cells<br>-40.8(± 9.4)% in female endocardial cells | [1] |
| NCX1 (SLC8A1) expression (%) | Human | No significant difference, ~10% larger in epicardial cells | [1] |
| | Human | +81(± 30)% larger at the base of the LV than the apex in non-menopausal females, not observed in males or post-menopausal females; NCX1 upregulated via estrogen. | [8] |
| | Rabbit, Rat | Upregulated NCX protein, mRNA expression and NCX activity in rabbit and rats. | [9] |
| $Na^+/K^+$-ATPase α1 expression (%) | Human | +84.5(± 27.6)% in female epicardial cells,<br>+64.8(± 22.4)% in female endocardial cells. | [1] |
| $Na^+/K^+$-ATPase α3 expression (%) | Human | -30.5(± 8.9)% in female epicardial cells<br>-30.4(± 7.8)% in female endocardial cells. | [1] |
| KCNA5 ($K_V1.5$) expression (%) | Human | No significant male-female differences. | [1] |
| RYR2 expression (%) | Human | No significant male-female differences. | [1] |

| | | | |
|---|---|---|---|
| SERCA2 ($Ca^{2+}$ ATPase 2) expression (%) Table 1 | Human | No significant male-female differences. | [1] |
| CALM3 expression (%) | Human | 23.5($\pm$ 6.9)% in female epicardial cells<br>27.2($\pm$ 6.7)% in female endocardial cells | [1] |
| PLN (Phospholamban) expression % | Human | -40.5($\pm$ 8.2)% in female epicardial cells<br>-24.7($\pm$ 12.4)% in female endocardial cells | [1] |

**Table S1**: Genomic expression data for male and female ion channel and transporter subunits taken into consideration for the sex-specific electrophysiological model.

In a similar approach to Peirlinck et al. (2021) [7], the male endocardial celltype was modified first, scaling directly from the ToR-ORd-Land baseline, and then used as a basis to scale other cell types and sex-specific differences for simplicity, with the additional benefit of being simple to implement into the Alya solver.

In addition to modifications from genomic expression data, minor adjustments were made to the RyR $Ca^{2+}$ release flux (reduced by 1%), SERCA2a $Ca^{2+}$ uptake rate (reduced by 3%), L-type $Ca^{2+}$ (increased by 5%) and $k_{off,tcl}$ was updated from the ToR-ORd model to address an issue in which the mid-myocardial cell model produced a spike-and-dome CaT morphology under conditions which increased intracellular $Ca^{2+}$ amplitude.

The conductance values for the relevant currents are presented in Table S2. Biomarkers for the three cell types are presented in the main text in Table 1; action potential, calcium transient and active tension traces are presented in Figure S1. The single cell simulated outputs for human ventricular male and female cells are presented in Table S3.

| **Parameter** | Male Endocardial (Baseline) Values |
|---|---|
| $G_{NaL}$ | 0.01774 |
| $G_{to}$ | 0.32 |
| $G_{pCa}$ | 0.00035 |
| $G_{Kr}$ | 0.03849 |
| $G_{Ks}$ | 0.00126 |
| $G_{K1}$ | 0.73318 |
| $G_{NaCa}$ | 0.00378 |
| $G_{NaK}$ | 13.9058 |
| $G_{Kb}$ | 0.01418 |
| $[Ca^{2+}]_{rel}$ | *0.99 |
| $[Ca^{2+}]_{up}$ | *0.94 |
| $CMDN_{max}$ | 0.04922 |
| $G_{CaL}$ | *1.05 |
| $k_{off,tcl}$ | 0.5 |

**Table S2: Male endocardial cell type current conductance values and scalings.** Values denoted by * represent a direct scaling applied to the final value obtained as in [2], due to the absence of a linear scaling factor available prior to calculation.

| | Female | | | Male | | |
|---|---|---|---|---|---|---|
| | **Endo** | **Epi** | **Mid** | **Endo** | **Epi** | **Mid** |
| $APD_{90}$ (ms) | 298 | 295 | 378 | 254 | 247 | 343 |
| $APD_{50}$ (ms) | 234 | 237 | 312 | 201 | 200 | 274 |
| Systolic $Ca^{2+}$ Peak (nM) | 454 | 689 | 898 | 515 | 776 | 944 |
| Diastolic $Ca^{2+}$ (nM) | 73 | 68 | 89 | 74 | 68 | 70 |
| $CaT_{90}$ | 368 | 323 | 381 | 343 | 297 | 356 |
| Peak Tension (kPa) | 22.6 | 40.5 | 60.4 | 27.6 | 45.2 | 62.5 |

**Table S3 – Single Cell Simulation Outputs for Human Ventricular Male and Female Celltypes**.
$APD_{90/50}$: time to 90/50% repolarisation of membrane potential. $CaT_{90}$: time to 90% reabsorption of systolic $Ca^{2+}$. Pacing = 1hz.

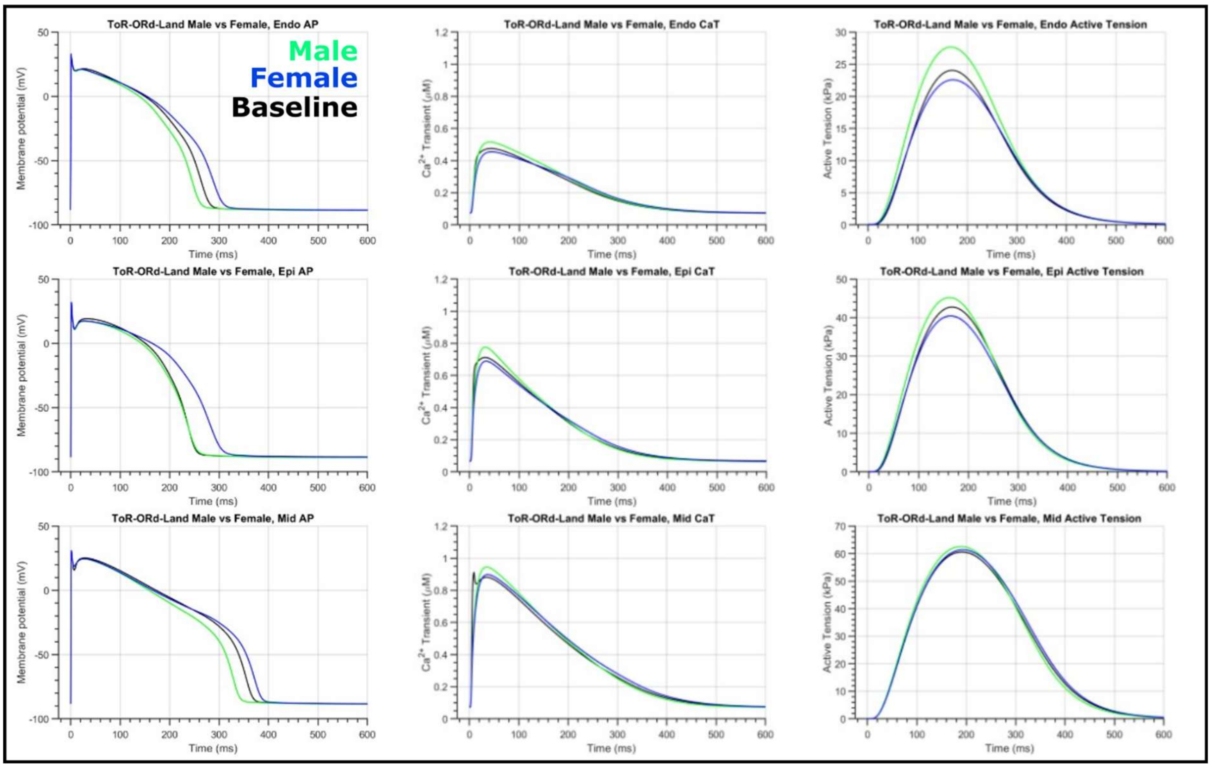

***SM Figure 1: Simulated sex differences in transmural ventricular cardiomyocyte behaviour.***
Male (green) and female (blue) human ventricular cell simulation outputs at steady state following 200 beats compared to baseline ToR-ORd-Land (black) simulation outputs. Final beat shown.

Simulations using these sex-specific variants were compared against the baseline model (Figure S1) to ensure deviations were within physiological ranges (Figure 2 in main text). In addition to this, trends in repolarisation, calcium transient decay and peak intracellular $Ca^{2+}$ over different pacing rates (see SM Figure 2) were also investigated and compared with sex-stratified data from the literature. $APD_{90}$ restitution dependence for each sex closely matches with previous sex-specific studies [10]. Simulated transmural $Ca^{2+}$ transient duration follows the same morphology, with similar trends in male and female (longer in female) presented in Fischer et al. (2016) [11] and peak intracellular $Ca^{2+}$ recapitulates experimentally observed differences at pacing rates from 0.5 Hz to 2Hz [12, 13].

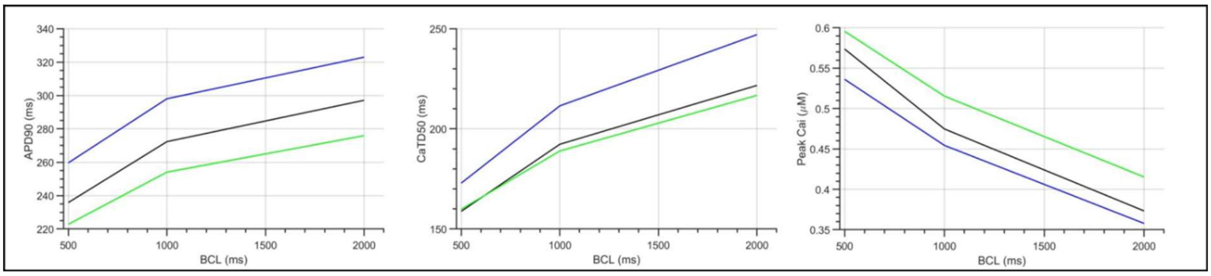

***Figure S2: Simulated trends in transmural ventricular cardiomyocyte behaviour at different pacing rates.***
Male (green) and female (blue) human ventricular cell simulation outputs represent biomarkers obtained from steady state following 200 beats at each pacing rate from 0.5 Hz to 2 Hz.

#### SM2. Electrophysiological Model and ECG Calibration

Electrophysiological heterogeneities in cardiac tissue are incorporated into our biventricular models as described in Wang et al. (2021) [14]. Transmural, apex-to-base and interventricular heterogeneities are included using experimental and clinical data. The exception to this rule is in the mid-myocardial cell type. Due to the lower accuracy of sex-differences and the minor difference the inclusion of mid-myocardial cells made at baseline; the mid-myocardial cell type is excluded from the biventricular simulations. The heterogeneities are modelled as two layers (endocardial, 70% of transmural width; epicardial, 30% of transmural width) with different properties as shown in Figure S1. The monodomain equation was used to simulate electrical propagation in the myocardium, and the same pseudo-ECG approach was utilised as in [14].

Activation patterns are inferred from a 12-lead QRS ECG segment using the methods found in Camps and Berg et al. (2024). Briefly, each root node has an activation time which is prescribed using a human model of the Purkinje tree network. The conduction speeds along the fibre, sheet-normal and Purkinje which best-match the clinical ECG are selected from an inferred population and these parameters are fixed to infer repolarisation characteristics.

A physiological apex-to-base electrophysiological heterogeneity was applied in both meshes through GKs scaling calibrated to clinical ECG t-wave morphology using a sequential Monte-Carlo approximate Bayesian computational inference method to infer repolarisation characteristics, as described in full in previous work [15]. Briefly, spatial variations in ionic currents underpinning repolarisation heterogeneity have been documented with the transient outward potassium current and the slow delayed rectifier potassium current ($I_{Ks}$) having robust evidence for spatial heterogeneity in the transmural, apex-to-base and transventricular directions. Of these, $G_{Ks}$ is chosen to be modulated because it had the larger effect on APD. $G_{Ks}$ is uniformly sampled between 1/50 and 50-fold its baseline value in ToR-ORd-Land.

The sequential Monte Carlo approximate Bayesian computation algorithm (SMC-ABC) is used to infer activation and repolarisation parameters from a clinical 12-lead ECG.

This method iteratively samples parameters from the 12-lead ECG and compares the resulting simulations until the population converges to one which complies with a specified cut-off discrepancy. This is used to determine a set of parameters which determine the ST segment and T-wave ECG signals. Outputs of this process are used to calibrate orthotropic conductivity parameters in the monodomain model to achieve desired conduction velocities.

#### SM3. Description of Mechanical Boundary Conditions

Mechanical boundary conditions are implemented as presented in the supplementary material of Wang et al. (2021) [14]; a short summary is given here along with linear parameters used for the male and female biventricular simulations.

Intra-ventricular pressure (P) and volume (V) behaviour is controlled independently in the left and right ventricles via a five-phased cardiac cycle state machine with a one-off initialisation phase. Both ventricles are inflated to a uniform endocardial pressure of 0.5kPa to reach a loaded resting endocardial volume. Following this one-time *initialisation* phase, both ventricles undergo *active inflation* in which the pressure in both ventricular chambers is linearly increased to an end diastolic pressure of 1.5 kPa over 100ms ($t_{diastole}$), mimicking the atrial contraction phase of diastolic filling. *Isovolumetric contraction* then occurs where the stimulated myocytes begin to generate active tension, and the endocardial pressure (P) is allowed to increase such that chamber volume is kept approximately constant. *Ejection* is triggered when the ventricular pressure exceeds the arterial pressure, set to be 9 kPa and 2 kPa in the left and right ventricles respectively. A two-element Windkessel model is used to simulate blood pressure of the systemic and pulmonary circulation systems. *Isovolumetric relaxation* is triggered by the reversal of ventricular volume change (i.e. $\frac{dV}{dT} > 0$), where pressure is allowed to decrease while volume remains approximately constant. Following relaxation, *passive filling* occurs in which the myocyte active tension is allowed to return to resting state, and the volume returns to the initialised rest value.

The parameters to reproduce the healthy baseline model are presented in SM Table 3. Input files for the simulations are available at **[Link will be provided for publication]**.

| Name | Parameter | LV | RV | Unit |
|---|---|---|---|---|
| Pericardial stiffness | $K_{epi}$ | 12185 | | Ba $cm^{-1}$ |
| Time to initial pressure | $\mathbf{t}_0$ | 0.02 | | s |
| Initial pressure | $P_0$ | 5000 | | Ba |
| Duration of passive diastolic filling | $t_{diastole}$ | 0.1 | | s |
| Pressure at end of diastole | $P_{endd}$ | 15000 | | Ba |
| Arterial compliance | C | 0.00075 | 0.002 | $cm^3$ $Ba^{-1}$ |
| Arterial resistance | R | 250 | 150 | Ba s $cm^{-3}$ |
| Aortic pressure | $P_{art0}$ | 90000 | 20000 | Ba |
| Pressure at end of isovolumetric relaxation | $P_{post}$ | 10000 | | Ba |
| Penalty parameters for isovolumetric contraction | $C_v$ | 0.3 | | $cm^3$ $s^{-1}$ $Ba^{-1}$ |
| Penalty parameters for isovolumetric relaxation | $C_v$ | 0.1 | 0.2 | $cm^3$ $s^{-1}$ $Ba^{-1}$ |
| Penalty parameters for passive filling | $C_p$, $C_v$ | 0.1, 1 | 0.2, 1 | $cm^3$ $Ba^{-1}$, $cm^3$ $s^{-1}$ $Ba^{-1}$ |

***Table S4: Passive mechanical parameters for boundary conditions and phase control.*** The units for pressure/stiffness is the Barye (Ba) which is consistent with the centimetre-gram-second system of units used in the simulations. 1 Barye = 10000 kPa.

#### SM4. Model Convergence

Electromechanics were simulated for five beats of 1000ms cycle length, achieving ECG and pressure-volume convergence after the second beat. Figure S3 demonstrates the ability of the model to return to resting state at the end of each beat, and convergence in these behaviours.

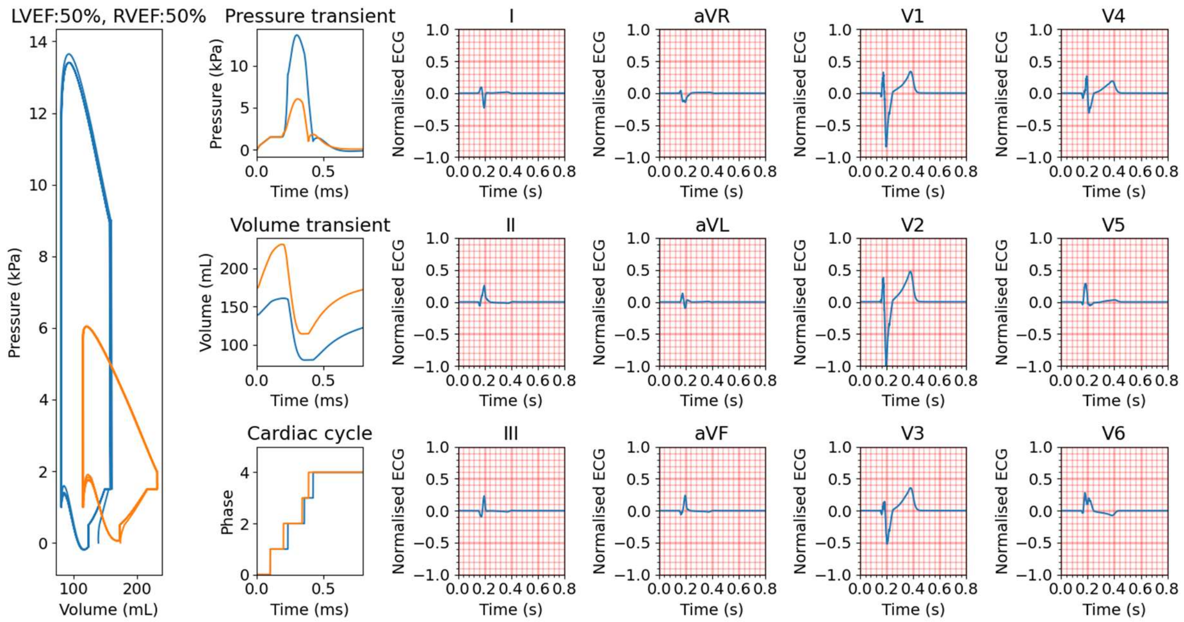

***Figure S3: Convergence of electromechanical simulations***

Five cycles of a healthy simulation showing pressure volume loop (left), pressure transient, volume transient, cardiac cycle phase transitions, and the whole ECG. Convergence was achieved from the second cycle onwards.

#### SM5. Models of Pharmacological Compounds

Drug effects were modelled using a simple pore block model, in which ionic conductances are multiplied by a scaling factor calculated using the following:

$$G_x = \frac{1}{1 + \left(\frac{[Drug\ Concentration]}{IC_{50}}\right)^h}$$

where $IC_{50}$ is the concentration at which the current is blocked by 50%, and the Hill coefficient exponent $h$ describes the gradient of block. Table S5 provides input data for four compounds used in this study for validation:

- Dofetilide, a class III antiarrhythmic drug which selectively blocks the hERG channel, used to maintain normal heart rhythm and cardioversion.

- Verapamil, a calcium channel blocker used to treat high blood pressure, and arrhythmogenic disorders. It primarily blocks L-type $Ca^{2+}$ channels, and secondarily acts as a hERG blocker.
- Ranolazine, an anti-anginal medication, which is used to improve blood flow. It primarily blocks the late sodium current, $I_{NaL}$ but also affects sodium, calcium and potassium currents.
- Quinidine, a class Ia anti-arrhythmic drug used to treat cardiac arrhythmias by restoring normal sinus rhythm. It inhibits potassium, calcium and sodium currents.

| Compound | $I_{Kr}$ $IC_{50}$ (nH) | $I_{CaL}$ $IC_{50}$ (nH) | $I_{Na}$ $IC_{50}$ (nH) | $I_{NaL}$ $IC_{50}$ (nH) | Source |
|---|---|---|---|---|---|
| Dofetilide | 0.001 (0.6) | - | - | - | [16] |
| Verapamil | 0.499 (1.1) | 0.202 (1.1) | - | - | [16] |
| Ranolazine | 10.9 (0.9) | 172 (0.6) | 30.2 (0.8) | 5.9 (1) | [17, 18] |
| Quinidine | 0.72(1.06) | 0.64(0.64) | 14.6 (1.22) | - | [19] |

***Table S5 – Drug Models used for simulating pharmacological effects.***

#### SM6. Repolarisation and Depolarisation Maps

To separately assess the effects of sex-specific electrophysiology from anatomical differences, biventricular simulations were performed on the male and female anatomies with all three human ventricular cell models. The female anatomy, when simulated with the female electrophysiology presented a smaller t-wave amplitude, a longer QTc interval and a shallower ST segment than when simulated with the baseline non-specific or male-specific cell models. Similarly, the male anatomy, produced the same effects when simulated with the female electrophysiology, compared with the non-specific and male-specific models.

We expected no observable change in the initial depolarisation of the anatomy with either sex-specific model, which held true in our simulations (Figure S4). Repolarisation was expected to be affected across the anatomies, primarily in late-stage depolarisation where this is determined by the balance of inward potassium channels and L-type $Ca^{2+}$ channel flux. We observed an approximate 20ms difference in repolarisation time across the female anatomy between the female and non-specific model, and symmetrically a small decrease in repolarisation time between the male and non-specific ionic models in the male anatomy (~ 10ms). This is well within expectations of repolarisation time differences between adult human males and females [20].

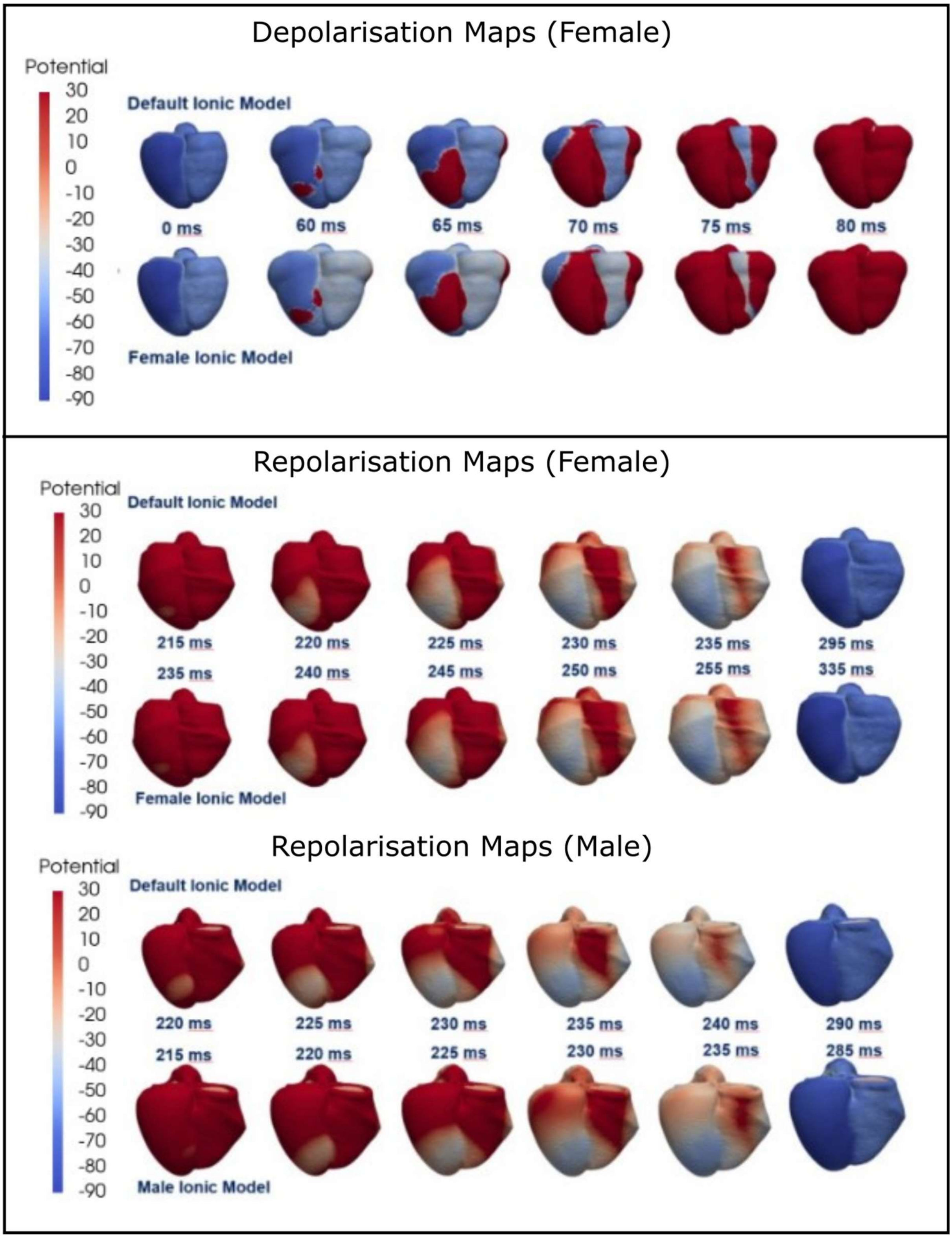

***Figure S3: Simulated Depolarisation and Repolarisation Maps.***
Electrical activity maps for the male and female anatomies, using the male- and female-specific ionic model variants presented in SM Figure 1, compared with the baseline non-specific model.

#### SM7. Sensitivity Analysis of Key Ionic Targets

The present study was conducted using state-of-the-art electrophysiological and mechanical models which were constructed based on human experimental and clinical data at the cellular and organ levels [4, 5]. Due to the tractability of the study, we rely on the validation performed in previous publications and here present a short analysis of the effects of variability in the key ionic drug targets presented in this paper on our two simulated outcomes of interest, QTc (Figure S5) and t-wave amplitude (Figure S6) for the male and female electrophysiological model. Conductance scaling factors for each current were analysed from scales of 10% to 100%, encompassing the range of drug blocks in the present study. $G_{Kr}$, $G_{CaL}$ and $G_{NaL}$ are the three primary blocks explored in this study, and $G_{Ks}$ is included due to its role in the ECG personalisation approach.

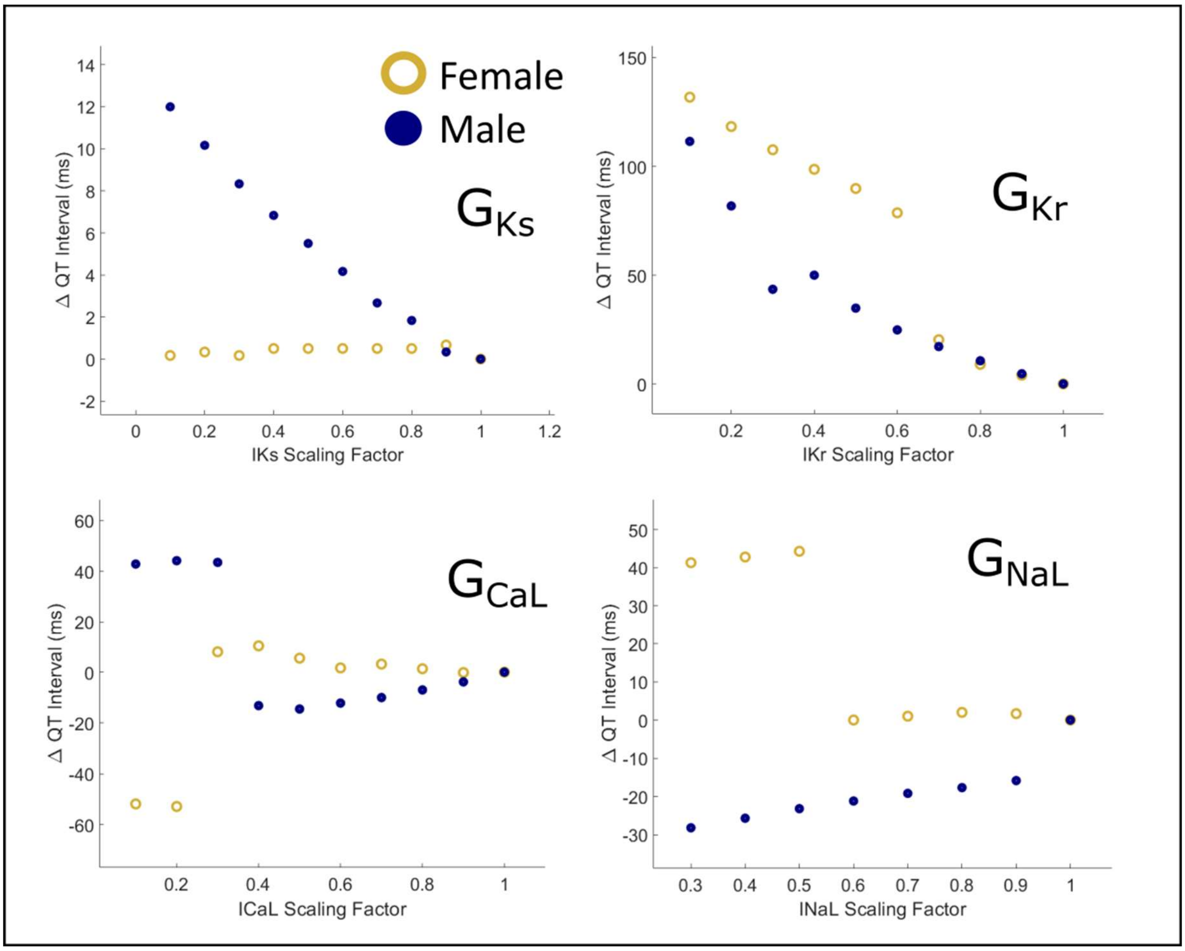

***Figure S4: Effects of Modifying Ionic Conductance on Simulated Male and Female QT interval***

Reductions of $G_{Ks}$ result in no change in QT interval in the female model, and a minor prolongation effect in the male model. $G_{Kr}$ block results in a much larger effect in both sexes, presenting similar prolongation up to 30% current block, and at blocks exceeding that, a notably larger effect in the female simulations; notably the female $G_{Kr}$ is reduced in comparison with the male as described in Table 1 of the main text. $G_{CaL}$ shows minor divergent impacts on simulated male and female QT prolongation, reflective of the relative balances of $G_{CaL}$ and inward potassium channels ($G_{Kr}, G_{Ks}$); these effects remain minor up to 60% block which exceeds any $Ca^{2+}$ block in the in-silico investigation. $G_{NaL}$ block results in no changes to simulated female QTc up to 40% block, which far exceeds the range of block for Dofetilide and Verapamil; but is relevant for Ranolazine and Quinidine which was not included in this study at the patient level. In the male simulations, $G_{NaL}$ block produced a linear dose-dependent minor decrease in QTc. To summarise, $G_{Kr}$ block has the largest pronounced impact on the QT interval in both male and female models, which is more pronounced in the female model due to reduced repolarisation reserves. At the drug concentrations implemented in this study, all other channel blocks will have minor to negligible effects QT duration.

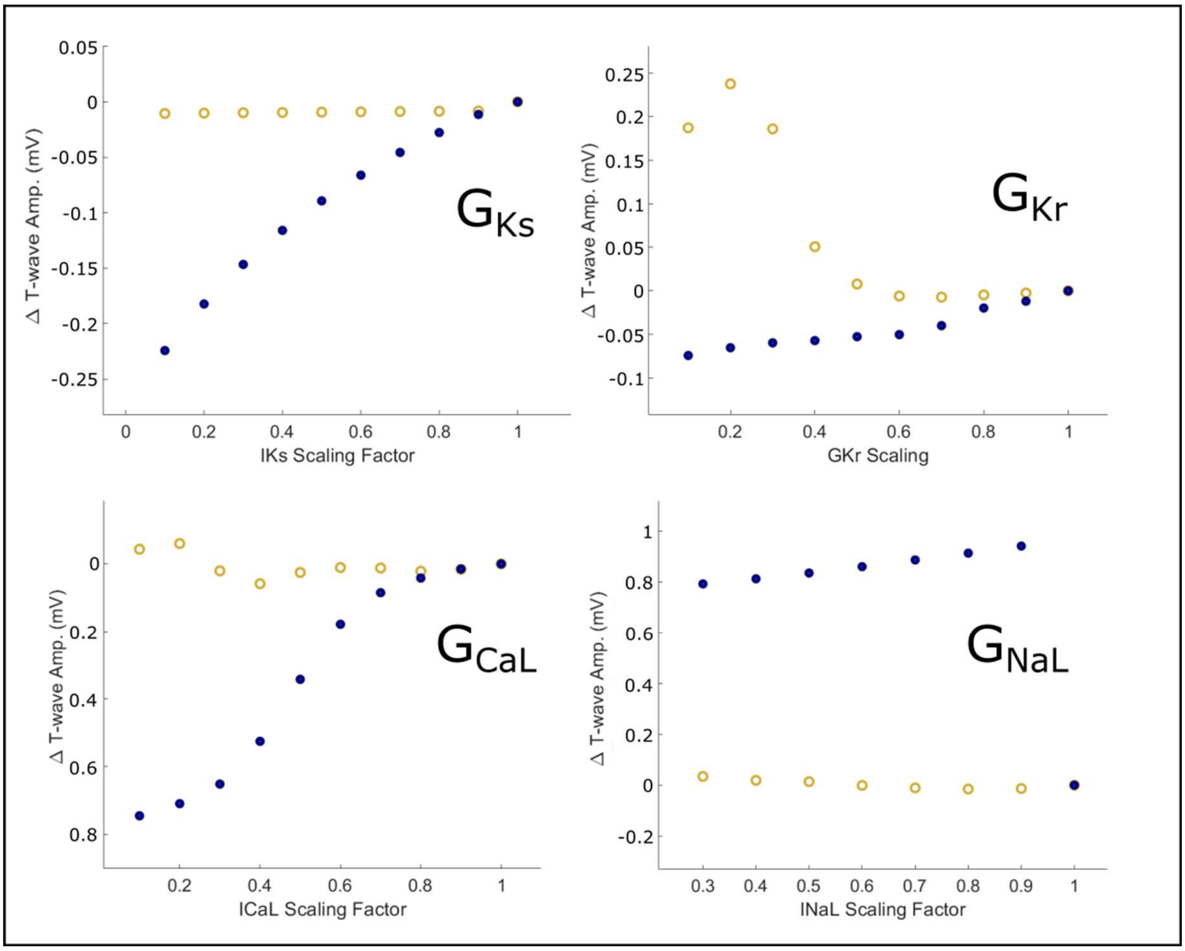

***Figure S6: Effects of Modifying Ionic Conductance on Simulated Male and Female t-wave amplitude.***

$G_{Ks}$ block has no effect on simulated female T-wave amplitude and a linear dose-dependent reduction in amplitude in the simulated male t-wave. $G_{Kr}$ block presents similar minor reductions in t-wave amplitude for both male and female simulations up to 30% block in which the trend bifurcates, with a continuing linear reduction in amplitude in the male simulations, and the female then showing an increase in t-wave amplitude. The t-wave describes ventricular repolarisation, thus the relative balances of phase 3 currents ($G_{CaL}$ vs. $G_{Kr}, G_{Ks}$, etc.) in ventricular myocytes collectively play a role in t-wave morphology. As we have only a single endocardial and epicardial cell model present, the impact of this balance may be more exaggerated than in vivo; heterogeneous cells will dampen this effect [21]. $G_{CaL}$ block has no effect on either male or female simulated t-waves up to 20% block, after which the male simulations demonstrate a dose-dependent reduction in t-wave amplitude up to 0.8 mV. $G_{NaL}$ shows no effect in the female simulations, and a consistent increase which is only minorly effected by dose-dependence in the male simulations.

#### SM8. Effect of ECG Personalisation Approach on ECG Outcomes

The ECG was calibrated using a sequential Monte-Carlo approximate Bayesian computational inference approach described in Camps et al. (2024) [22]. Electrical activation and repolarisation characteristics are inferred from a clinical 12-lead ECG, with fast simulations performed using a reaction-Eikonal model. $G_{Ks}$ scaling is utilised to vary cellular APD throughout the anatomies to produce a close match of the simulated ECG and clinical ECG selected. To achieve this through $G_{Ks}$ scaling alone, the baseline ionic model is modified by scaling the conductance of $G_{Kr}$ by 70%, $G_{Ks}$ five-fold, and decreasing the time-constant of L-type $Ca^{2+}$ channel activation from 75ms to 60ms. As demonstrated in SM6, this will have impact on the simulated ECG, thus this effect was tested in non-specific simulations under dofetilide by enabling and disabling the ECG personalisation (SM Figure 6). Inclusion of the ECG personalisation changes resulted in approximately a 110 ms increase simulated QT for all doses of dofetilide tested. Additionally, simulated t-wave amplitude showed a divergence from minor decreases in the simulated t-waves to minor increases. This behaviour may have played a role in the over-estimation of simulated drug effects in this study.

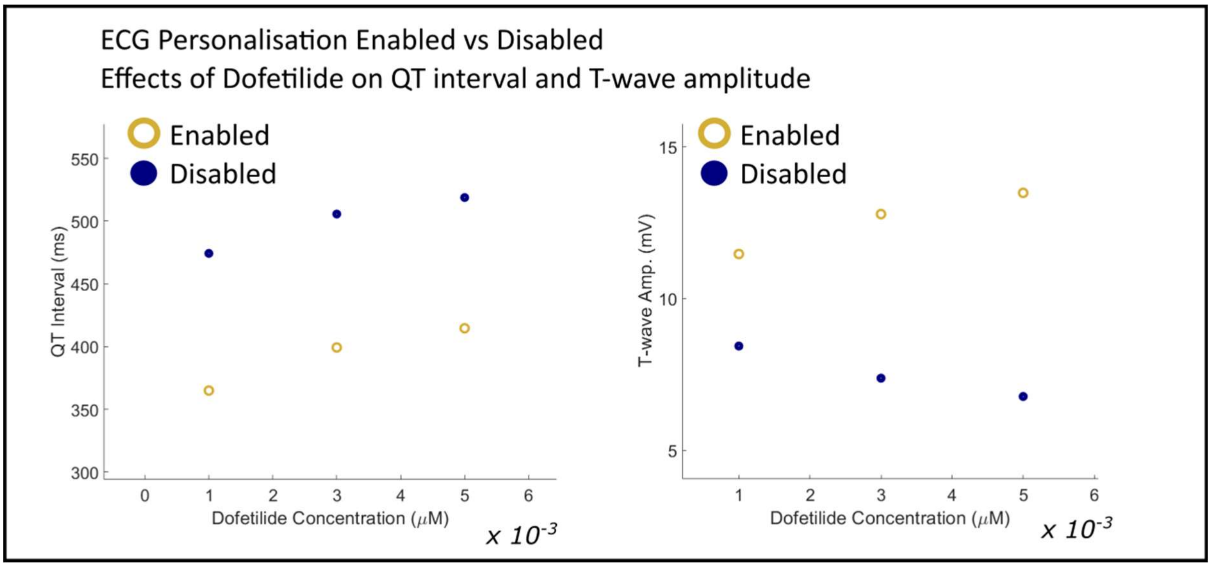

**Figure S7: Effect on QT interval and T-wave amplitude of ECG personalisation methodology.**

#### SM9. Effect of changing passive mechanical parameters on ECG.

The effects of modulating mechanical parameters in the contraction model on the ECG were investigated to assess the impact of potential sex-specific differences in mechanics would result in significant changes to the ECG (Figure SM7). The parameters tested are $T_{ref}$ scaling, $K_{ws}$ scaling, pericardial stiffness $K_{epi}$, bulk modulus Kct, arterial compliance $C$, arterial resistance $R$, gain relaxation $C_v$, and passive mechanical terms $a, as, asf$ in the Holzapfel-Ogden model [23]. Each of these were scaled from 50-200% of the baseline value.

No mechanical parameters tested were observed to have any significant impact on the resulting simulated ECG, and thus we can surmise that the ECG changes observed in this study on each anatomy by sex-specific modulation or drug modulation are a result of electrophysiological changes only.

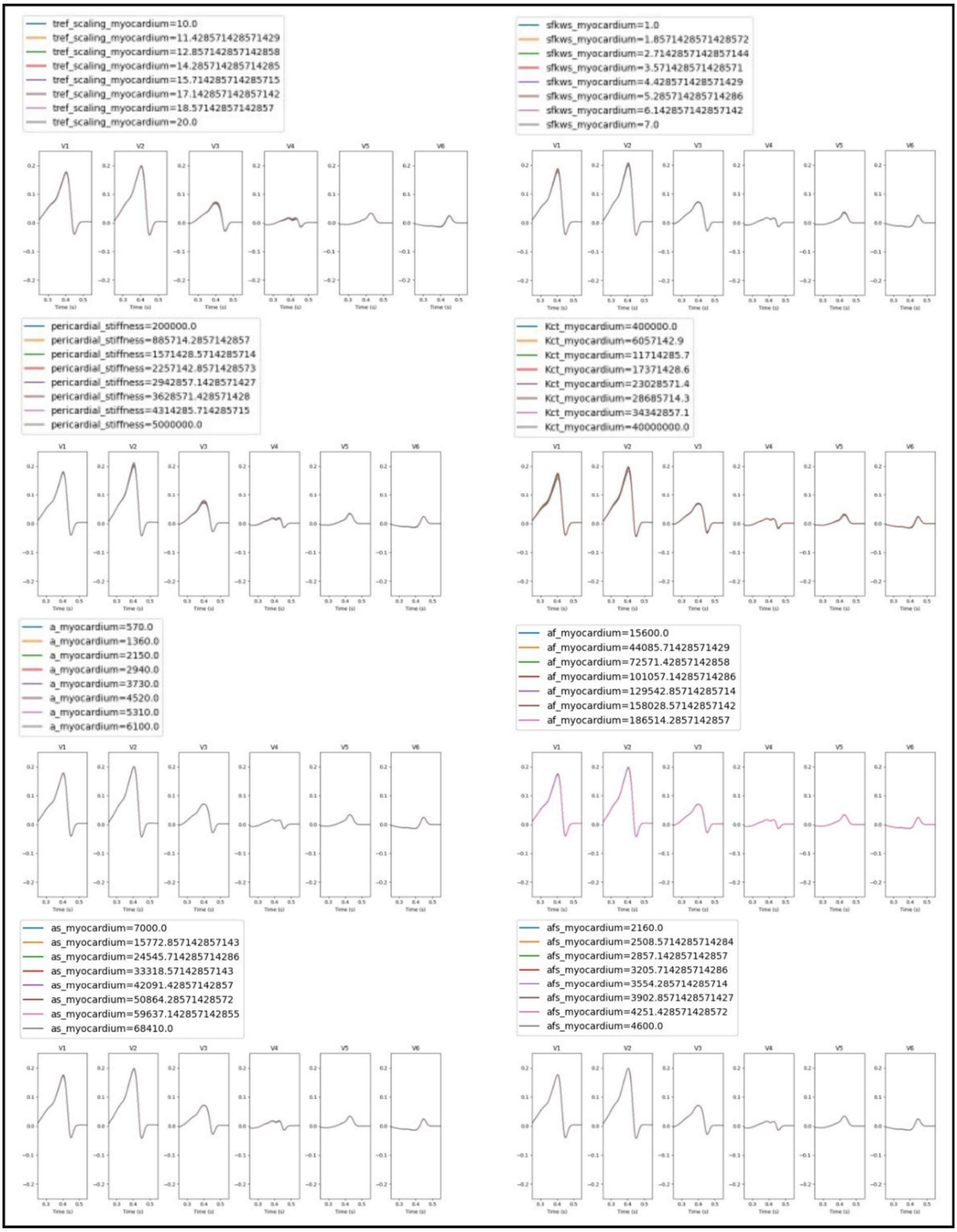

***Figure S8: Effects of modulating mechanical parameters on the ECG.***
One beat after convergence in healthy non-specific model shown overlayed for each value in legend above showing effects on all pre-cordial leads.
